# Polycomb sustains promoters in a deep OFF-state by limiting PIC formation to counteract transcription

**DOI:** 10.1101/2023.06.13.544762

**Authors:** Aleksander T. Szczurek, Emilia Dimitrova, Jessica R. Kelley, Robert J. Klose

## Abstract

The Polycomb system plays fundamental roles in regulating gene expression during mammalian development. However, how it controls transcription to enable gene repression has remained enigmatic. Here we employ rapid degron-based depletion coupled with live-cell transcription imaging and single-particle tracking to uncover how the Polycomb system controls transcription in single cells. We discover that the Polycomb system is not a constitutive block to transcription but instead sustains a long-lived deep promoter OFF-state which limits the frequency with which the promoter can enter into a transcribing state. We demonstrate that Polycomb sustains this deep promoter OFF-state by counteracting the binding of factors that enable early transcription pre-initiation complex formation and show that this is necessary for gene repression. Together these important discoveries provide a new rationale for how the Polycomb system controls transcription and suggests a universal mechanism that could enable the Polycomb system to constrain transcription across diverse cellular contexts.

## Introduction

The capacity to initiate and maintain defined gene expression patterns is fundamental to complex multicellular development. At its most basic level, this relies on transcription factors recognizing DNA sequences in gene regulatory elements to control RNA polymerase II (RNA Pol II) activity at the core gene promoter^1^. However, in eukaryotes, chromatin states at gene regulatory elements can also profoundly influence transcription and gene expression, and the systems that create these states are essential for normal gene regulation and development^1–4^. While there is an emerging appreciation of the mechanisms through which transcription factors instruct transcription^1^, how chromatin-based systems influence transcription remains very poorly understood and a major conceptual gap in our knowledge of gene regulation.

The Polycomb repressive system represents a paradigm for chromatin-based gene regulation and is essential for appropriate gene expression during animal development^5–7^. It comprises two distinct histone modifying complexes, Polycomb repressive complex 1, and 2 (PRC1 and PRC2, respectively). PRC1 mono-ubiquitylates H2A at lysine 119 (H2AK119ub1) and PRC2 methylates histone H3 at lysine 27 (H3K27me3). In vertebrates, both PRC1 and PRC2 are targeted to promoters of genes that have CpG island elements. Here they can deposit histone modifications and through feedback mechanisms create Polycomb chromatin domains that have high levels of H2AK119ub1, H3K27me3, and occupancy of PRC1/2 complexes^6^. Polycomb chromatin domains play important roles in counteracting gene expression and help to maintain the inactive state of genes in tissues where they should not be expressed^5–7^, with recent work also suggesting a more pervasive role in constraining gene expression^8–12^. However, how the Polycomb system controls transcription to repress gene expression remains very poorly understood.

A central experimental constraint that has limited our understanding of how gene regulatory mechanisms function in situ is that the process of transcription is not uniform across cells. Instead, transcription is stochastic within individual cells over time and varies substantially between cells in a population^13, 14^. As such, ensemble approaches for analyzing transcription do not capture key features of the transcription cycle that are essential for understanding how regulatory mechanisms achieve their effects on gene expression. To overcome this limitation, single-cell transcription analysis complemented with detailed understanding of the cellular dynamics of the factors that regulate transcription is emerging as an important new avenue to uncover how transcription is controlled to regulate gene expression^13, 14^.

We and others have recently demonstrated using ensemble approaches in embryonic stem cells (ESCs) that the Polycomb system, in particular PRC1 and H2AK119ub1^11, 15–20^, plays a central role in constraining gene expression through limiting the activity of RNA Pol II at its target genes^21^. This has demonstrated that the factors necessary to promote transcription of Polycomb target genes are present and that the Polycomb system must limit some key aspect of transcription to enable repression. Further analysis of these effects in single cells suggested that the Polycomb system could influence the frequency of transcriptional bursts, but this observation relied on inferring kinetic parameters based on modelling RNA-transcript levels in fixed cells^21–23^. As such, how the Polycomb system controls transcription remains essentially unknown.

To address this fundamental question, here we exploit rapid degron approaches, live-cell imaging, and genomics to uncover how PRC1/H2AK119ub1 regulates transcription. We discover that PRC1/H2AK119ub1 plays an important role in sustaining a deep promoter OFF-state by limiting transcription pre-initiation complex (PIC) engagement with gene promoters to counteract transcription. As such, we reveal that Polycomb chromatin domains limit the earliest steps of transcription to enable gene repression.

## Results

### Imaging Polycomb target gene transcription in live cells

To begin understanding how the Polycomb system influences transcription we employed a highly sensitive MS2 aptamer-based system which is capable of capturing transcription with single-transcript sensitivity in living cells (Figure 1A)^24^. To implement this, we used CRISPR-Cas9 engineering in mouse ESCs to create lines where MS2 repeats were inserted into the first intron of genes that have promoters which are embedded within Polycomb chromatin domains and whose expression are counteracted by the Polycomb system (*Zic2* and *E2f6*) and a moderately expressed reference gene that lacks a discernable Polycomb chromatin domain (*Hspg2*) (Supplementary Figure S1A, B, C). These cell lines were also engineered to express MS2 RNA binding protein fused to GFP (MCP-GFP), allowing nascent transcription imaging and quantification of transcription in live cells^24^.

**Figure 1.**
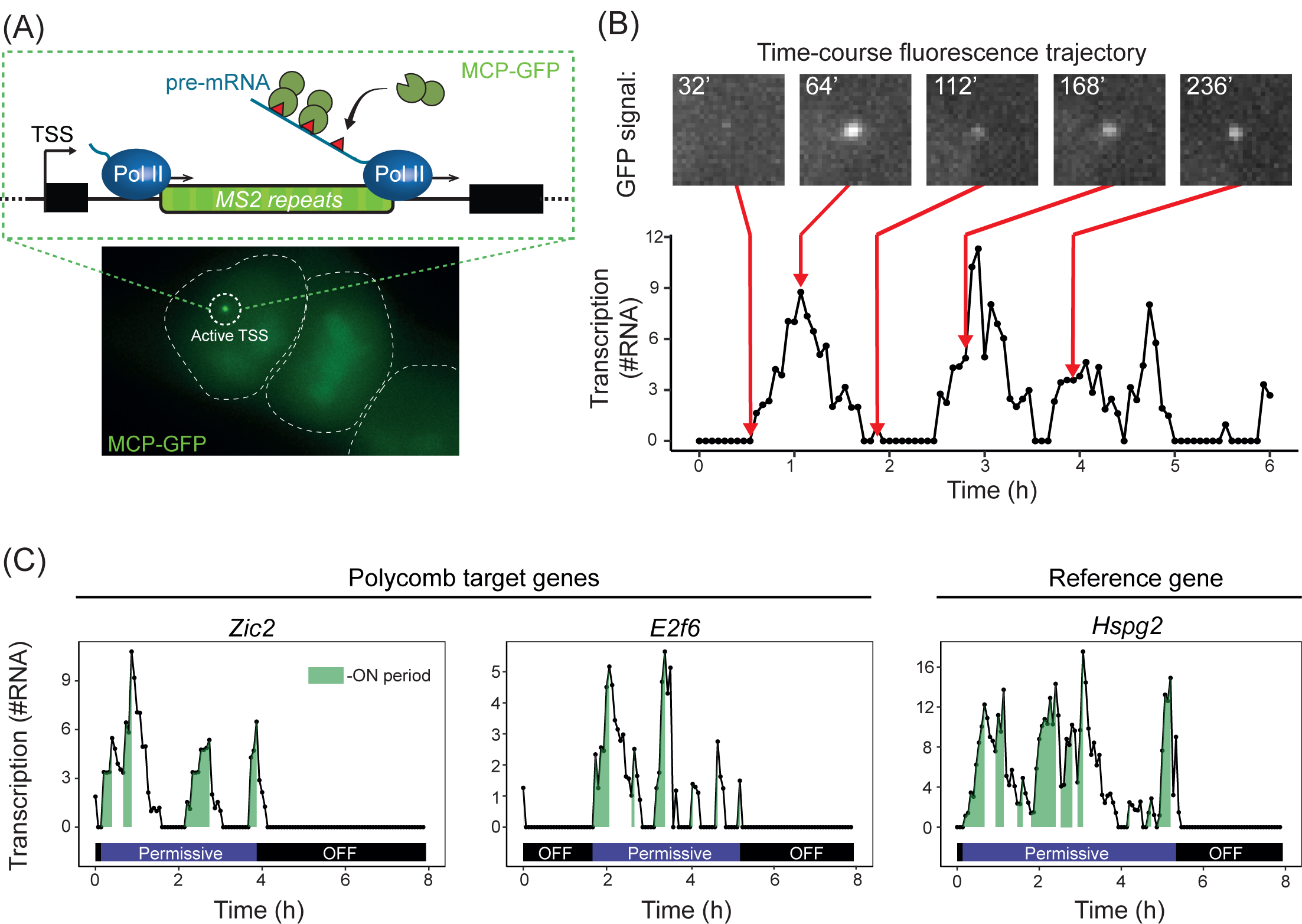
Imaging Polycomb target gene transcription in live cells. A) A schematic illustrating the transcription imaging approach. MS2 repeats were inserted into a promoter-proximal intron of the genes of interest. As RNA Pol II passes through the array, nascent RNA presents MS2 stem loops that are bound by MCP-GFP leading to accumulation of fluorescence signal at the active transcription site (top). Bottom: an example image of a cell with a nascent transcription spot corresponding to active transcription start site (TSS). The white dashed lines indicate the cell outline. B) An example of a transcription activity trajectory from cells engineered to contain the MS2/MCP-GFP system (*Zic2)*. Maximal projections of the focalized MCP-GFP signal are shown above the trajectory to illustrate the pulsatile nature of transcription. C) Example transcription activity trajectories for Polycomb target genes (*Zic2* and *E2f6*) and a reference gene (*Hspg2*). ON (green), permissive (violet), and OFF-periods (black) are illustrated. The Y axis represents transcriptional activity (in RNA molecules).

When we imaged these cell lines, bright MCP-GFP foci were evident which corresponded to nascent RNA-FISH signal for each gene (Supplementary Figure S1B), and we found that nascent transcription could be quantified in live cells with single transcript sensitivity (Supplementary Figure S3C, D). Importantly, transcription of Polycomb target genes was detected in agreement with these genes being expressed, albeit at low levels. When we measured MCP-GFP fluorescence signal corresponding to nascent transcription over time for each gene we observed that transcription was pulsatile (Figure 1B), in line with previous live-cell transcription imaging in mammalian cells^13, 14^. Furthermore, transcription trajectories for all three genes were characterized by what appeared to be transcriptionally permissive-periods within which there were distinct bursts of transcription initiation, that we refer to as ON-periods, where multiple RNA polymerases transcribe in close succession (Figure 1C). Permissive-periods were interspersed by long-lived OFF-periods during which the gene was not transcribed at all. Some of these OFF-periods were highly persistent, extending for the entire duration (8 hrs) of the imaging movie, and clonal expression analysis demonstrated that in some instances these OFF-periods could extend across cell divisions (Supplementary Figure S2A, B)^24, 25^. Therefore, our imaging approach captures the transcriptional behaviour of Polycomb target genes, and provides us with an opportunity to study how the Polycomb system regulates transcription in live cells.

### PRC1 does not constrain transcription during ON-periods

With the capacity to image the transcription of Polycomb target genes we could begin to explore how the Polycomb system might regulate transcription. Initially we focused on ON-periods and developed a transcription imaging analysis approach that allowed us to extract the number of transcripts, the duration, and the PolII loading frequency during ON-periods (Figure 2A, Supplementary Figure S3E, F). When we compared ON-period features for Polycomb-target genes (*Zic2* and *E2f6*) and the reference gene (*Hspg2*) we were surprised to find that they were similar (Figure 2B) despite Polycomb genes being much more lowly expressed (Figure 2D).

**Figure 2.**
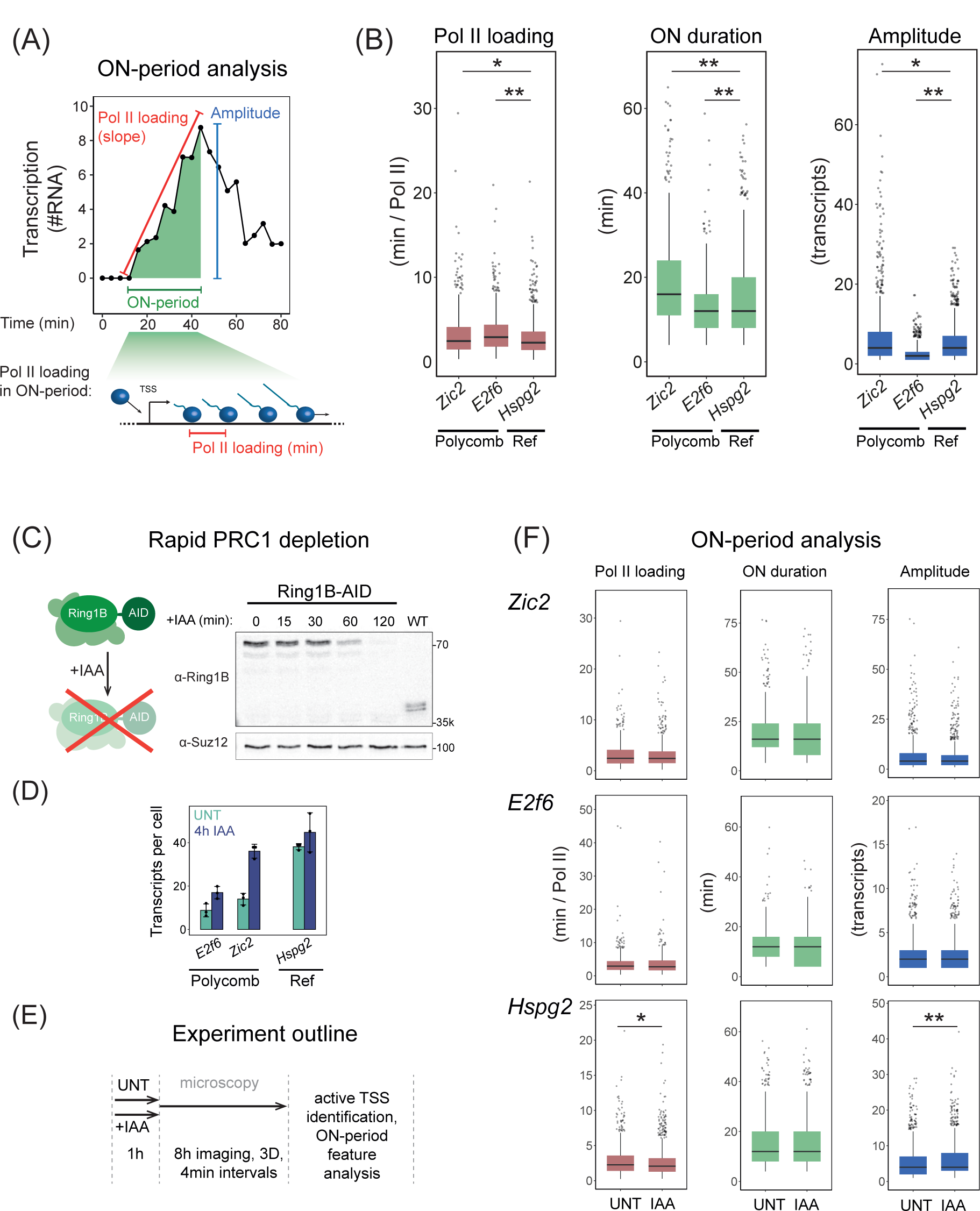
PRC1 does not constrain transcription during ON-periods. A) A schematic illustrating the ON-period features extracted from transcription imaging trajectories. These include the rate of RNA Pol II initiation within the ON-period (from linear fit of the slope), the duration of the ON-period (min), and the amplitude of the ON-period (transcripts). B) Box plots comparing the ON-period features showing the median values, interquartile range (IQR), whiskers as 1.5 IQR, and outliers as dots. P-values were estimated using a Kolmogorov-Smirnov (KS) test with <0.05 (indicated as *) and <0.01 (indicated as **). Box plots represent data from 4, 3, 2 biological replicates for *Zic2, E2f6*, and *Hspg2*, respectively. C) A diagram illustrating the auxin inducible system used to rapidly deplete the catalytical subunit of PRC1 (RING1B) (left). Western blot analysis of Ring1B-AID levels over a 2-hour period after addition of auxin (IAA) (right) compared to a WT mESC line. D) Analyses of *E2f6*, *Zic2*, and *Hspg2* expression 4h after PRC1 depletion using smRNA-FISH. The dots represent individual biological replicates (n=3, with >400 cells per replicate) and error bars represent the standard deviation. E) A schematic illustrating the approach to image transcription in live cells with (IAA) or without (UNT) PRC1 depletion. F) Box plots corresponding to ON-period analysis for *Zic2*, *E2f6,* and *Hspg2* (Ref) in untreated (UNT) and PRC1-depleted (IAA) conditions.

This suggested that Polycomb-mediated repression may not primarily manifest from limiting transcription during ON-periods. To test this, the MS2 reporter system was integrated into a degron cell line where addition of the small molecule auxin (IAA) leads to rapid depletion of the catalytic subunit of PRC1 and turnover of H2AK119ub1 (Figure 2C)^21, 26^. Importantly, PRC1/H2AK119ub1 depletion caused Polycomb target gene derepression and resulted in a roughly 2-2.5 fold increase in transcript levels as assessed by single molecule RNA-FISH (smRNA-FISH), with *Zic2* reaching transcript levels similar to the reference gene (Figure 2D). We then examined ON-period features and found they were largely unaffected after PRC1/H2AK119ub1 depletion despite these genes displaying elevated transcript levels (Figure 2E, F). Therefore, we conclude that Polycomb-mediated repression is not achieved by PRC1 constraining transcription during ON-periods.

### PRC1 sustains a deep OFF-state that is refractory to transcription

PRC1/H2AK119ub1 depletion did not cause major effects on transcription during ON-periods suggesting that PRC1/H2AK119ub1 must regulate some other feature of transcription to repress gene expression. One possibility was that PRC1 could limit the frequency of transcription events (ON-periods) during permissive-periods, or the duration of permissive-periods (Figure 3A). To test this possibility, we imaged transcription in the presence or absence of PRC1 and quantified the time between ON-periods within permissive-periods (Figure 3B), and the duration of permissive-periods (Figure 3C). Similarly to transcription ON-period analysis, this revealed that for Polycomb target genes depletion of PRC1/H2AK119ub1 had only minor effects on transcription during permissive periods, although we did see a small increase in the duration of permissive-periods for the reference gene. Therefore, PRC1/H2AK119ub1 does not appear to repress Polycomb target genes via regulating either ON-period (Figure 2) or permissive-period features (Figure 3B, C).

**Figure 3.**
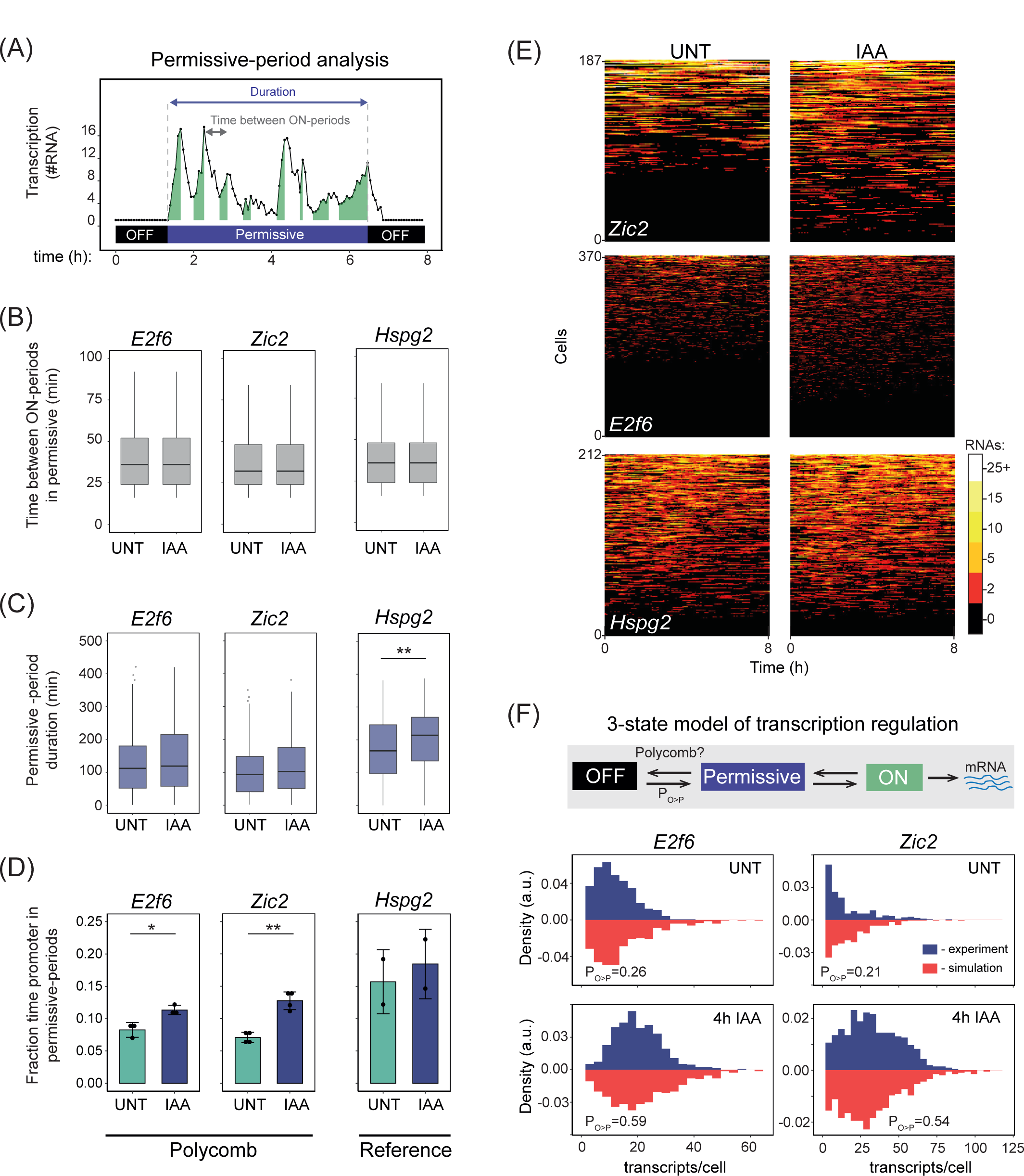
PRC1 sustains a deep OFF-state that is refractory to transcription and counteracts gene expression. A) A schematic illustrating the features extracted from transcription imaging trajectories for permissive-period analysis. These include the time between ON-periods within permissive-periods (grey arrow) and the duration of permissive-periods (purple arrow). B) Box plots comparing the time between ON-periods for *E2f6*, *Zic2*, and *Hspg2* in untreated (UNT) or PRC1-depleted (IAA) conditions showing IQR, median, and whiskers as 1.5 IQR. Throughout the figure * or ** are shown if the KS p-value was <0.05 or <0.01. C) Box plots comparing the duration of permissive-periods for *E2f6*, *Zic2*, and *Hspg2* in untreated (UNT) or PRC1-depleted (IAA) conditions. D) Bar graphs showing the fraction of total imaging time spent in permissive-periods for *E2f6*, *Zic2*, and *Hspg2.* Bars correspond to mean values, error bars standard deviation, and dots are biological replicates (4, 3, 2 for *Zic2, E2f6*, and *Hspg2*, respectively). E) Heatmaps illustrating transcription imaging trajectories of individual cells for *E2f6*, *Zic2*, and *Hspg2* in untreated (UNT) or PRC1-depleted (IAA) conditions over the 8-hour imaging time course (horizontal-axis). The amplitude of transcription is illustrated in the scale bar (right) and the number of imaging time courses is indicated on the y-axis. Heatmaps were randomly subsampled to represent equal number of measurements in UNT and IAA to facilitate qualitative comparison. F) A schematic illustrating the simple 3-state model of transcription used to simulate gene expression distributions (top). Histograms comparing transcript per cell distributions from smRNA-FISH in experiments (blue bars, experimental) and simulations (red bars) for Polycomb target genes in untreated (UNT) or PRC1-depleted (IAA) conditions. The best-fit p_O>P_ value for both UNT and 4h IAA are indicated.

Having observed little effect of PRC1/H2AK119ub1 on either ON-period or permissive-period features, we postulated that the effects on expression must instead manifest from an increase in the frequency with which Polycomb target genes exit from long-lived OFF-periods and enter into permissive-periods where transcription occurs. Consistent with this possibility, when we examine the fraction of time that promoters spend in permissive-periods, we discovered that depletion of PRC1/H2AK119ub1 caused a clear increase, despite permissive period duration remaining largely unaltered (Figure 3D). This increase in the frequency of permissive-periods was also evident in heatmaps illustrating single-cell imaging traces for Polycomb target genes (Figure 3E). Therefore, PRC1/H2AK119ub1 counteracts transcription by sustaining promoters in a long-lived (Supplementary Figure S2) deep OFF-state.

### Derepression of Polycomb target genes corresponds to an increased probability of exiting the deep OFF-state

If PRC1/H2AK119ub1 represses transcription by sustaining a deep OFF-state, it would follow that an increased frequency of transitioning out of this deep OFF-state should account for the elevated gene expression observed in smRNA-FISH following PRC1/H2AK119ub1 depletion (Figure 2D). To investigate this possibility, we built a simple three state gene expression model that incorporated parameters measured in live-cell imaging for ON-periods (Figure 2B), the number of ON-periods and time between them within permissive-periods (Supplementary Figure S4A, B, C, D), and transcript half-lives (Supplementary Figure S4E). Stochastic simulations of gene expression were then carried out with differing probabilities of transitioning from OFF-periods to permissive-periods (p_O>P_, Figure 3F) to identify p_O>P_ values that corresponded to the transcript distributions measured by smRNA-FISH in untreated cells (Supplementary Figure S4G). We then asked whether simply increasing the p_O>P_ value in these gene expression simulations would reproduce the increased expression and transcript distributions measured in cells when PRC1/H2AK119ub1 was depleted (Figure 2D, 3F). Importantly, for both *E2f6* and *Zic2* this demonstrated that an approximately 2.5 fold increase in p_O>P_ resulted in similar transcript distributions to those observed experimentally after PRC1/H2AK119ub1 depletion, consistent with this being the point of transcriptional control (Figure 3F). Therefore, by combining live-cell imaging, stochastic simulations, and gene expression analysis we show that the Polycomb system sustains a long-lived deep promoter OFF-state that is refractory to transcription to repress gene expression.

### Single particle tracking reveals that PRC1 counteracts binding of early PIC forming components

The process of transcription is orchestrated by a number of distinct regulatory mechanisms that contribute to transcript production^1, 27, 28^. To understand how PRC1 sustains the deep OFF-state, we set out to define what regulatory feature of transcription PRC1/H2AK119ub1 controls. The behavior of individual factors that regulate the core process of transcription are, like the process of transcription itself, known to be stochastic and highly dynamic. Therefore, capturing the breadth of their dynamic behaviors is not possible using classical ensemble genomic approaches. However, it has recently been shown that these dynamic behaviors can be measured and quantified in living-cells using single particle tracking (SPT), where the dynamics of individual molecules can be directly observed as they interact with binding sites on chromatin^29–35^. Therefore, we reasoned that similar approaches could be applied to explore the regulatory stage of transcription affected by PRC1/H2AK119ub1.

To enable single particle tracking we used CRISPR-Cas9 genome engineering to endogenously HALO-tag a series of core transcription regulators that represent distinct steps in transcription^27, 28^ (Figure 4C and Supplementary Figure S5A, B). To examine early transcription initiation, we HALO-tagged the TATA-box binding protein (TBP), and the TAF1 and TAF11 components of TFIID^36^. TBP function in pre-initiation complex (PIC) formation can be counteracted by the negative cofactor 2 (NC2) which binds to a surface on TBP that is required for engagement of the general transcription factors TFIIA and B^37^. Therefore, we HALO-tagged NC2β to capture inhibition of early PIC formation and TFIIB whose interaction with TBP is essential for progression of PIC formation^38^. PIC formation then progresses through binding of the Mediator coactivator complex, so we also HALO-tagged the core component of the Mediator complex MED14. Once RNA Pol II has engaged with the PIC, TFIIH is recruited through its contacts with Mediator and RNA Pol II^39, 40^ and its CDK7 component phosphorylates the C-terminal domain heptapeptide repeats of RNA Pol II during early transcription elongation. Therefore, we HALO-tagged CDK7 to capture this step of transcription. As RNA Pol II enters into early elongation CDK9 phosphorylates the negative elongation factor (NELF) and RNA Pol II to overcome RNA Pol II pausing and ensure productive transcription elongation. To capture factors related to this stage of transcription we HALO-tagged CDK9, NELF-B, and the largest subunit of RNA Pol II, RPB1.

**Figure 4.**
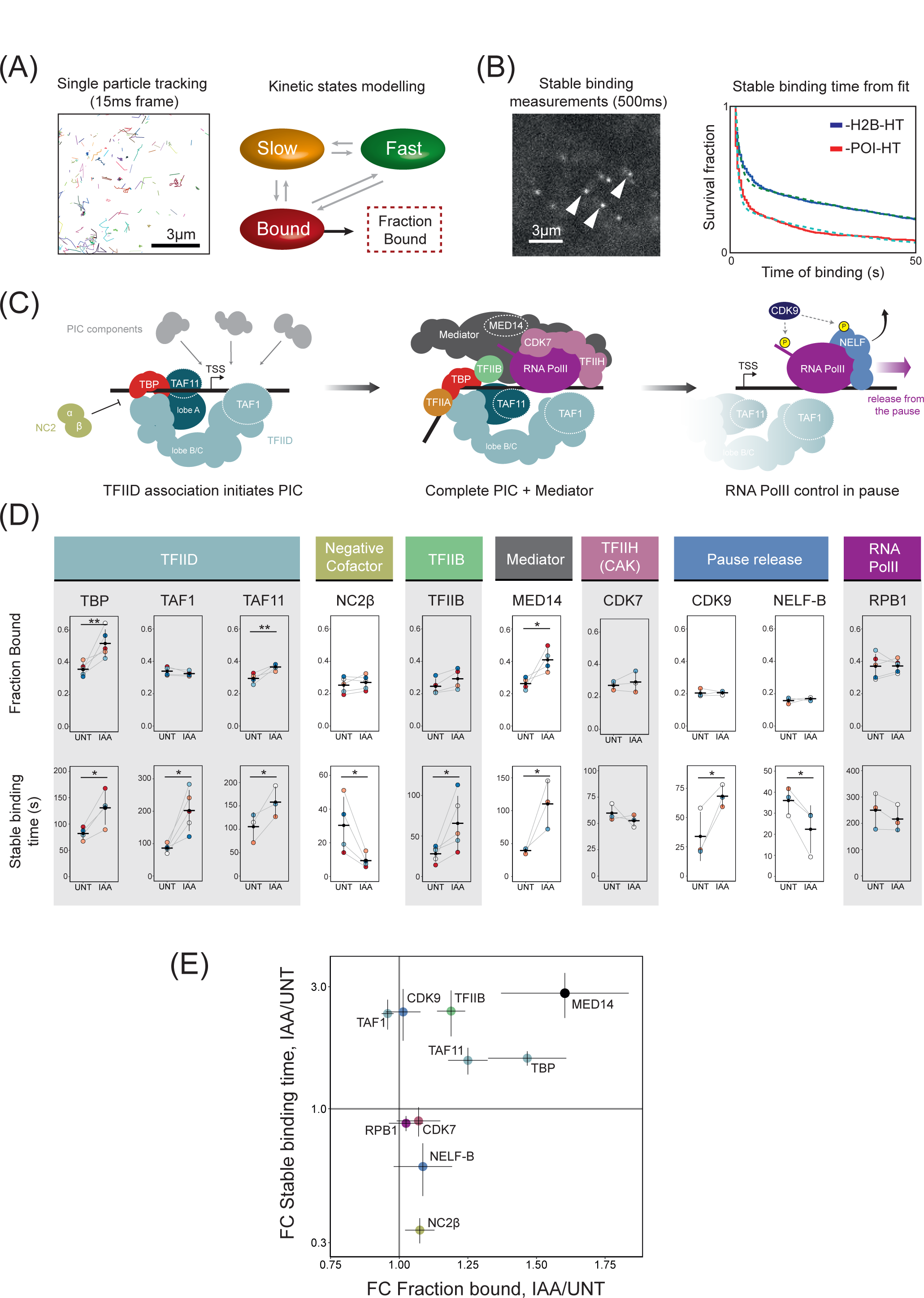
PRC1 counteracts binding of early PIC forming components. A) An example of individually color-coded single molecule tracks acquired at high frame rate (left). These tracks are used for kinetic modeling in SPOT-ON^42^ to obtain bound factions. B) An example frame from stable binding time measurements acquired at low frame rate with stably bound molecules indicated with arrow heads. Stable binding times for the protein of interest (POI) are extracted from bi-exponential fits (dotted lines) from cumulative distributions (solid lines) and corrected for photobleaching using estimates of stable binding of histone H2B-HT (blue). C) A cartoon illustrating stages of PIC assembly and transcription regulation. Protein factors studied by SPT are indicated. D) Dot plots illustrating the bound fractions (top) and stable binding time (bottom) for a panel of transcription regulators in untreated (UNT) or PRC1-depleted (IAA) conditions. Individually color-coded dots represent values for individual biological replicates and are connected with grey lines, error bars represent standard deviation, and horizontal lines show the mean value. P-values represent one-sided t-test with * indicating p-values of <0.05 and ** indicating <0.01. A Minimum of 3 biological replicates were measured with approximately 100 cells per replicate for bound fraction analysis and approximately 20 cells for stable binding time measurements per biological replicate. E) A scatter plot integrating the effects on bound fraction and stable biding times measured in SPT. Dots correspond to the mean fold change (FC) values for individual proteins and the error bars correspond to standard error of the mean. Solid grey vertical and horizontal lines correspond to 1 (no change).

To image the behavior of these transcription regulators in single-cells with single-molecule precision we employed a photoactivatable HALO dye coupled with highly inclined and laminated optical sheet (HILO) microscopy^41^. We carried out imaging at a high frame rate to quantify the fraction of molecules bound to chromatin (association, Figure 4A and Supplementary Figure S5C)^42^ and also carried out imaging at a low frame rate to estimate the stable binding time of molecules (dissociation, Figure 4B and Supplementary Figure S5D)^43^. Interestingly, by focusing on the earliest regulatory steps involving TBP (Figure 4C), we observed that PRC1/H2AK119ub1 depletion resulted in a nearly 50% increase in the bound fraction of TBP and its binding time also increased (Figure 4D). This indicates that TBP engages more frequently and remains bound for longer in the absence of PRC1/H2AK119ub1. When we examined the dynamics of other TFIID components, TAF11 showed an increased bound fraction whereas TAF1 was unaffected, but both factors displayed increases in stable binding time. It has been proposed that the A lobe of TFIID, which contains TAF11, and the B/C lobe of TFIID, which contains TAF1, may exist in distinct preassembled subcomplexes^44, 45^. Our analysis therefore suggests that PRC1/H2AK119ub1 may primarily influence engagement of TBP and TFIID lobe A, with the net result being more stable binding of the entire TFIID holocomplex. In contrast to effects on TFIID, the bound fraction of the TBP inhibitory factor NC2β was largely unaffected, but its duration of binding was dramatically reduced consistent with elevated stable binding of a TBP-containing TFIID complex. In agreement with the effects on TBP/TFIID, the bound fraction and duration of MED14 binding was also elevated upon PRC1 depletion consistent with recent evidence that Mediator engagement is dependent on TFIID^46^. This suggests that in the absence of PRC1/H2AK119ub1, the association and stable binding of early PIC forming components is increased, whereas the stable binding time of the negative cofactor complex is reduced.

To understand whether these early effects would also influence downstream general transcription factors, we next examined TFIIB and the TFIIH subunit CDK7 (Figure 4D). TFIIB showed only a slight increase in bound fraction but was subject to elevated stable binding, whereas CDK7 was largely unaffected. We then examined CDK9 and NELF-B and found that their bound fractions were unaffected, but the stable binding time of CDK9 increased whereas it decreased slightly for NELF-B, in line with elevated transcription initiation when PRC1/H2AK119ub1 is depleted^47^. Importantly, when we measured RNA Pol II via examining RPB1 dynamics we observed very little effect, supporting the idea that PRC1 regulates early transcription events and does not significantly affect the amount of elongating RNA Pol II which is what we primarily capture in our measurements. Furthermore, this result indicates that the increase in the amount of elongating RNA Pol II that occurs at more lowly expressed Polycomb target genes, does not contribute significantly enough to the overall amount of elongating RNA Pol II to influence our measurements. Based on these detailed kinetic measurements, we find that PRC1/H2AK119ub1 limits the binding of factors involved in the earliest stages of PIC formation (Figure 4E).

### PRC1 constrains TFIID binding to inhibit gene expression

Single particle tracking suggested that PRC1/H2AK119ub1 may counteract the binding of TFIID to limit the very earliest regulatory steps of transcription. While single particle tracking captures the breadth of transcription factor binding dynamics with single molecule precision, it is not capable of providing information about where the effects on binding occur in the genome. To understand where TFIID binding was affected we carried out calibrated chromatin-immunoprecipitation coupled to massively parallel sequencing (cChIP-seq) for endogenously tagged TAF1 before and after PRC1 depletion. We chose TAF1 as it is the largest subunit of TFIID and a component of the TFIID holocomplex^36^. Importantly, when we segregated Polycomb-enriched and non-Polycomb transcription start sites (TSSs) based on PRC1 occupancy, we observed the highest levels of TAF1 at non-Polycomb genes (Figure 5A) in line with these genes being more highly expressed. Importantly, we also observed TAF1 binding at Polycomb-enriched genes, but the levels were much lower, in line with the repressed state of these genes and consistent with the idea that PRC1 could limit TFIID complex binding to sustain a deep promoter OFF-state. To test this possibility, we depleted PRC1 and observed a clear increase in TAF1 occupancy at Polycomb enriched genes (Figure 5A, B). Interestingly, we also observed a modest yet significant increase in TAF1 binding across non-Polycomb enriched transcription start sites, indicating that PRC1 may constrain the binding of TFIID more broadly (Figure 5B, Supplementary Figure S6B). Consistent with this possibility, low levels of PRC1 are detected at non-Polycomb gene promoters, and when we analysed gene expression across these genes we also observed a modest increase in expression after PRC1 depletion (Supplementary Figure S6A). These findings are in line with previous observations that PRC1 and H2AK119ub1 may also have more subtle yet pervasive effects on gene expression^11, 21^. Together these observations are consistent with PRC1 limiting transcription and gene expression by counteracting TFIID binding to gene promoters, with the largest effects occurring at lowly transcribed Polycomb target genes that have high levels of PRC1 and H2AK119ub1.

**Figure 5.**
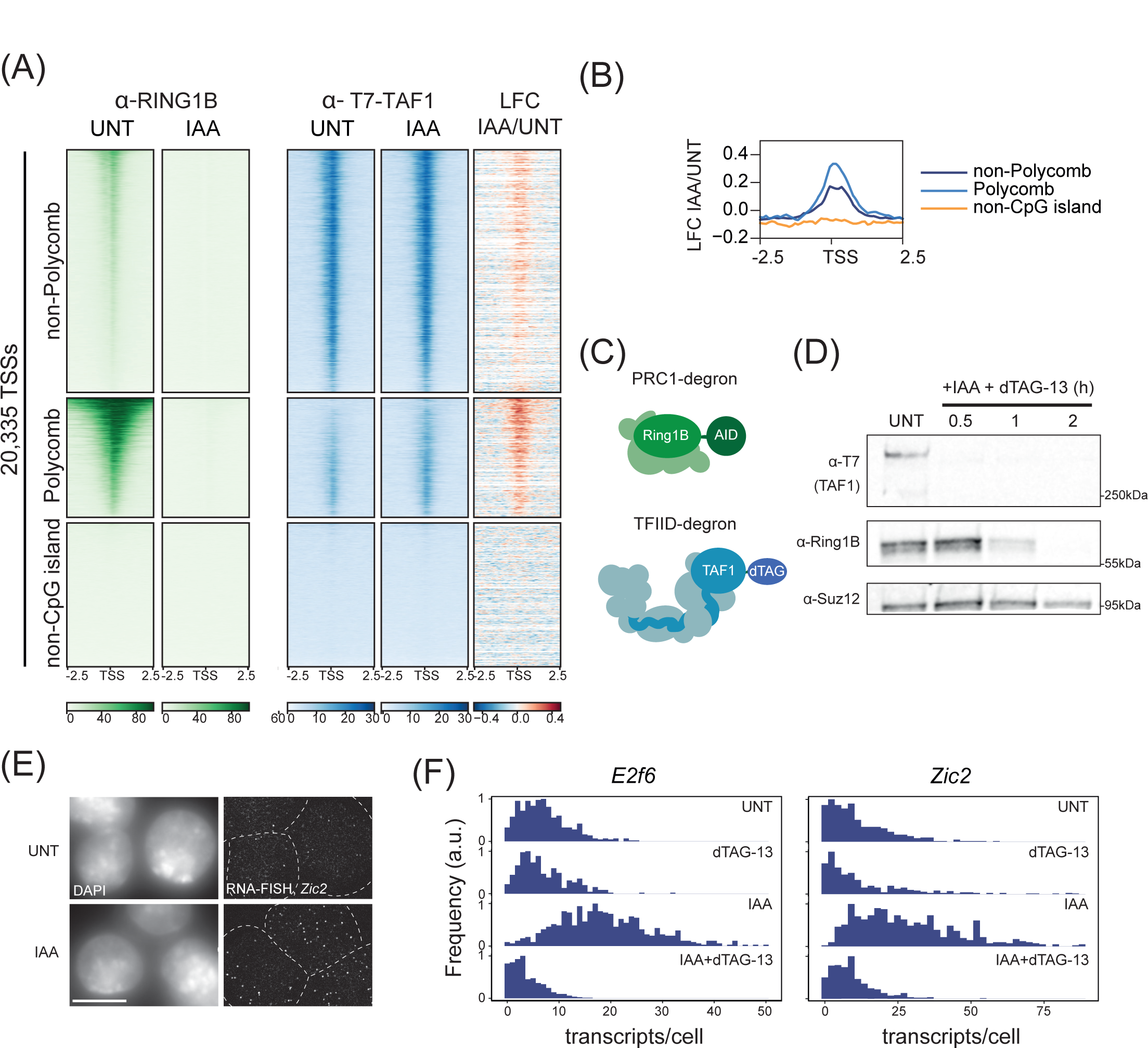
PRC1 constrains TFIID binding to inhibit gene expression. A) A heatmap illustrating cChIP-seq signal for RINGB (PRC1) (green, left) or endogenously T7-tagged TAF1 (blue, right) in untreated (UNT) or treated (IAA) ESCs across transcription start sites (TSS). The distance in kilobases from left and right of TSSs is shown below each heatmap. To visualise changes in T7-TAF1 signal, the Log2 fold change (LFC) IAA/UNT is shown to the right of the T7-TAF1 cChIP-seq signal. TSSs were segregated into non-Polycomb (n=9899), Polycomb (n=4869), and non-CpG islands (n=5869) groupings as indicated and ranked by RING1B signal. B) A metaplot illustrating the Log2FC IAA/UNT of T7-TAF1 cChIP-seq signal at the three classes of TSSs shown in A. C) A schematic illustrating the combinatorial degron strategy used to examine the contribution of TFIID to derepression of Polycomb target genes after depletion of PRC1. D) Western blot to analyse the levels of RING1B-AID and dTAG-TAF1 after simultaneous addition of IAA and dTAG-13 over a 2 h time course. SUZ12 is shown as a loading control. E) A smRNA-FISH image labeling *Zic2* (Polycomb target) transcripts in untreated (UNT) or after 4h of RING1B-depletion (IAA) illustrating increased transcript numbers. White dashed lines indicate cell outlines and scale bar represents 10 µm. F) smRNA-FISH analysis of transcript-per-cell distributions for untreated (UNT), cells with TAF1 depleted (dTAG-13), cells with RING1B-AID depleted (IAA), and cells with both RING1B and TAF1 depleted (IAA + dTAG-13). Depletions were performed for 4h and at least 400 cells were measured for each gene in each condition.

Given that PRC1/H2AK119ub1 depletion caused increased TFIID binding at Polycomb target genes and an increased propensity to exit from the deep transcriptional OFF-state, we wondered whether TFIID was required for the derepression of Polycomb target genes. Therefore, we used CRISPR-Cas9 to engineer a degron tag into the endogenous *Taf1* gene in the PRC1 degron cell line (Figure 5C, D) as the TAF1 protein is integral to the formation of the TFIID holocomplex^45^. We then depleted either PRC1 or PRC1 and TAF1 simultaneously and examined the expression of the *Zic2* and *E2f6* Polycomb target genes using smRNA-FISH (Figure 5E, F). Importantly, this revealed that neither Polycomb target gene was derepressed in the absence of TAF1, suggesting that TFIID binding enables elevated expression in the absence of PRC1/H2AK119ub1. Therefore, we discover that Polycomb-mediated gene repression relies on sustaining a deep OFF-state through limiting TFIID binding at gene promoters.

## Discussion

How chromatin states regulate transcription to control gene expression has remained a major conceptual gap in our understanding of gene regulation. Here we approach this fundamental question by focusing on the Polycomb system. Through exploiting rapid degron-based protein depletion, transcription-imaging, and simulations, we discover that the Polycomb system counteracts transcription by sustaining promoters in a long-lived deep OFF-state (Figure 1-3). By measuring how transcription regulatory factors interact with chromatin using live-cell single-particle tracking and genomic approaches, we demonstrate that the Polycomb system sustains this deep OFF-state by counteracting the binding of factors that enable early PIC formation (Figure 4). Finally, using degron approaches and gene expression analysis we demonstrate that Polycomb target gene derepression relies on increased association of TFIID, demonstrating an important role for the Polycomb system in limiting the association of general transcription factors (Figure 5). Together these discoveries provide a new rationale for how the Polycomb system regulates transcription.

A number of distinct models have previously been proposed to explain how the Polycomb system influences transcription to counteract gene expression^6, 21, 48–55^. However, these are mostly derived from *in vitro* biochemistry or ensemble fixed-cell analyses that is blind to the dynamic control processes that regulate transcription in living cells. Our live-cell transcription imaging now reveals that PRC1/H2AK119ub1 primarily functions to repress transcription and gene expression by limiting transition out of a deep promoter OFF-state and into the permissive-state where ON-periods or bursts of transcription occur. We demonstrate that this effect is mediated by counteracting association of early PIC components with the promoter, consistent with recent observations demonstrating that alterations in TATA box sequences that reduce their affinity for TBP and manipulating factors that affect PIC formation also limit entry into permissive-periods^22, 24, 56, 57^. Therefore, we identify central role for Polycomb-mediated and chromatin based gene repression in regulating the OFF-to-permissive promoter state transition.

Importantly, our new findings in live cells differ from previous *in vitro* biochemical observations suggesting that Polycomb complexes might block recruitment of Mediator, but not TBP/TFIID^48^. We believe this may be related to the fact that chromatin templates used in *in vitro* reconstitution experiments do not contain H2AK119ub1, which we and others have recently shown is important for repression *in vivo*^15, 16^. Unlike most other histone modifications, ubiquitylation is a bulky 76 amino acid adduct that dramatically alters the nucleosome, suggesting it could possibly function to repress transcription by influencing how transcription and other regulatory factors interact with promoter chromatin^39, 40^. Recent biochemical and structural work has shown that TFIID and other components of the general transcription machinery make key contacts with nucleosomes as part of early transcription initiation mechanisms^40^. With this in mind, an important avenue for future biochemical and structural work will be to understand whether H2AK119ub1 influences how the core transcriptional machinery interacts with promoter chromatin to enable gene repression.

Gene expression can be very dynamic throughout mammalian development. For example, genes may be inactive during early development and their repression maintained by the Polycomb system, but later in development their expression may be required. Consistent with this requirement, we now discover that Polycomb-dependent repression does not act as a constitutive block to transcription but instead functions by limiting binding of early PIC forming components to reduce the probability that a promoter enters into a transcriptionally permissive-state. Given the breadth of gene types that the Polycomb system must presumably regulate in distinct cellular contexts, limiting the function of general transcription factors may provide universal means to constrain transcription at genes with diverse regulatory inputs without having to influence highly divergent gene-specific DNA binding factors or other regulatory influences. In the context of developmental transitions when Polycomb target genes become activated, we envisage that limiting the frequency of entering into permissive-periods could also ensure low-level activation signals are quelled, yet the gene promoter would remain receptive to strong and persistent activation signals necessary to initiate gene expression. Interestingly, once genes are activated, persistent transcription leads to Polycomb chromatin domain erosion in part through the transcriptional machinery guiding Trithorax-chromatin modifying systems which install histone modifications that inhibit Polycomb chromatin domain integrity^5, 6, 58^. This suggests Polycomb and Trithorax systems may counteract each other by installing chromatin states that decrease or increase the probability that a gene promoter is in a state that is permissive to transcription. In the context of future work it will be important to uncover whether this control point is the focus of antagonistic Polycomb/Trithorax systems.

In conclusion, we demonstrate that the integration of rapid degron approaches, live-cell imaging of transcription, and detailed analysis of transcription regulatory factors by single particle tracking can provide new insight into how chromatin-based gene regulation is controlled in living cells. In doing so, we provide compelling new evidence that PRC1/H2AK119ub1 represses gene expression by sustaining promoters in a deep OFF-state that is refractory to PIC formation and transcription.

## Materials & Methods

### Cell culture

The Ring1A^-/-^, Ring1B-AID mouse embryonic cell line was previously described and extensively characterized^21, 26^. Cells were grown on gelatinised culture plate at 37C and 5% CO_2_ in medium containing Dulbecco’s Modified Eagle Medium (Gibco) with 10% foetal bovine serum (Sigma), 2 mM L-glutamine (Life Technologies), 1x non-essential amino acids (Life Technologies), supplemented with 0.5 mM β-mercaptoethanol (Life Technologies) and 10 ng/ml leukemia inhibitory factor (produced in house) and split every other day. To deplete Ring1B-AID the cells were treated with indole-3-acetic acid (auxin, Life Technologies) at 500 µM whereas to deplete T7-dTAG–TAF1 the cells were treated with 20 µM 5,6-Dichloro-1-beta-D-ribofuranosylbenzimidazole for 1h, washed three times, and treated with 100 nM dTAG-13 for 4h^59^ (Tocris).

### Genome engineering

In order to knock-in HALO-Tag^60^, FKBP12^F36V^ (dTAG, Addgene #62988), MS2×128 array^24^, or MCP-GFP (Addgene #40649) into specific genomic location (typically N- or C-termini of a gene, or the first intron for MS2 array) guide sequences were designed using CRISPOR tool^61^ and cloned into pSptCas9(BB)-2A-Puro(PX459)-V2.0 guide expression plasmid (Addgene #62988). The complete list of guide sequences can be found in Supplementary Table 1. Targeting constructs used as templates for homology-directed repair were Gibson-assembled using Gibson master-mix (New England Biolabs) and PCR-amplified homology arms corresponding to genomic sequence flanking the desired site of insertion. A list of primers used to amplify homology arms are included in Supplementary Table 2. MCP-GFP, dTAG, or HALO-Tag were PCR-amplified from respective plasmids. The MS2×128 array was cut out of its original plasmid (a kind gift from E. Bertrand)^24^ using AleI/NheI restriction enzymes. dTAG was Gibson assembled to include a 3xT7-3xStrep-tag. To carry out targeting cells were transfected with 2 µg of the targeting construct and 0.5µg of the guide expressing construct using Lipofectamine 3000 according to the manufacturer’s protocol (Thermo Fisher Scientific). 1 day after transfection cells were plated sparsely and selected with 1 µg/ml puromycin for 48h. Puromycin was removed and the cells were grown until distinct colonies formed. Individual clones were picked and propagated in 96 well plates that were then screened for homozygous insertion by PCR. Screening primers are available in Supplementary Table 2. HALO and dTAG tagging was validated at protein level by western blot and in the case of HALO-Tag by labelling it with tetramethylrhodamine (TMR) and microscopy (Supplementary Figure S5A, B). MCP-GFP cells were inspected for expression uniformity (Supplementary Figure S3A). The integrity of MS2×128-containing lines was further confirmed by PCR using Q5 (New England Biolabs) and Terra (Takara) polymerases as well as by microscopy using RNA-FISH detecting intronic sequences (Supplementary Figure S1B) expected to colocalize with nuclear MS2×128/MCP-GFP foci.

### Nuclear extraction and western blot

Nuclear extraction and western blot analysis were performed as described previously^21^. Briefly, for nuclear extraction, cells growing on 10cm plate were harvested, washed once with PBS, and resupsended in 10 volumes of Buffer A (10 mM HEPES pH 7.9, 1.5 mM MgCl2, 10 mM KCl, 0.5 mM DTT, 0.5 mM PMSF and protease inhibitor cocktail (Roche)). Subsequently, cells were spun down at 1500g for 5min, and resuspended in 3 volumes of buffer A with 0.1% NP-40. Following centrifugation, the pellet was resuspended in 1 volume of Buffer C (5 mM HEPES (pH 7.9), 26% glycerol, 1.5 mM MgCl 2, 0.2 mM EDTA, protease inhibitor cocktail (Roche) and 0.5 mM DTT) with 400 mM NaCl and incubated on ice for 1h. Nuclei were pelleted by centrifugation at 16,000g for 20min at 4C. The supernatant was retained as nuclear extract (NE). For western blotting, 15-20 µg of NE was heated in SDS loading buffer at 95C for 5min and loaded on to an 8-12% acrylamide gel or a tris-acrylamide NuPAGE gradient gel (Thermo Fisher Scientific) and separated by electrophoresis. Next, the resolved proteins were transferred onto nitrocellulose membrane using Trans-Blot Turbo Transfer System (Bio-Rad). The membrane was blocked with 5% milk in PBS/0.1% Tween-20 (PBST-milk) for 1h. The membrane was transferred to PBST-milk containing primary antibodies (Supplementary Table 3) and incubated overnight at 4C. The following day memberanes were washed 3x with PBST-milk and incubated for 1h with secondary antibody conjugated with IRDye (Li-COR). Following 3×5min washes with PBST and a 5min wash with PBS the membrane was visualised with Odyssey Fc system (Li-COR).

### Chromatin immunoprecipitation and high throughput sequencing

Calibrated chromatin immunoprecipitation (cCHIP) was performed as previously described^62^. In brief, 5×10^7^ ES cells engineered with T7-dTAG-TAF1 were fixed with 1% formaldehyde (methanol-free, Thermo Fisher Scientific) for 10 min at 25C under constant gentle rotation. Fixation was quenched with 150 mM glycine and the cells were washed with ice-cold PBS and snap frozen in LN2. Additionally, 5×10^7^ HEK293T T7-SCC1 cells (a gift from Martin Houlard) were fixed with 1% formaldehyde as above and snap frozen in 2×10^6^ aliquots.

For spike-in calibration, 2×10^6^ HEK293T cross-linked cells were resuspended in 100 µl ice-cold lysis buffer (50 mM HEPES pH 7.9, 150 mM NaCl, 2 mM EDTA, 0.5 mM EGTA, 0.5% NP-40, 0.1% sodium deoxycolate, 0.1% SDS) and added to 5×10^7^ fixed ESCs resuspended in 900 µl lysis buffer. The cells were incubated on ice for 10 minutes and sonicated using Bioruptor Pico sonicator (Diagenode) for 23 cycles (30s ON/30s OFF), shearing genomic DNA to produce fragments between 300bp and 1kb.

Prior to immunoprecipitation, chromatin was diluted to 300 µg/ml with lysis buffer and precleared with Protein A agarose beads (Repligen), blocked with BSA and tRNA, for 1 hour at 4C. The precleared chromatin was then incubated with 5µl of anti-T7 antibody (Cell signaling) overnight rotating at 4C. Antibody-bound chromatin was purified with 20µl blocked Protein A agarose beads for 3 hours at 4C. ChIP washes were performed as described previously^62^. ChIP DNA was eluted in 1%SDS and 100 mM NaHCO_3_ and reversed crosslinked at 65C with 200 mM NaCl and RNase A (Sigma) under constant shaking. The samples were then treated with 20 µg/ml Proteinase K (Sigma) and purified using a ChIP DNA Clean and concentrator kit (Zymo Research). The corresponding input DNA was purified for each sample. The efficiency of each ChIP reaction was confirmed by quantitative PCR.

For cChIPseq, three reactions were set up for each condition and pooled for library preparation. Prior to library preparation, 5 ng ChIP DNA was diluted to 50 µl in TLE buffer (10 mM Tris-HCl pH 8.0, 0.1 mM EDTA) and sonicated with a Bioruptor Pico sonicator for 17 min (30 s on and 30 s off). Libraries were prepared using NEBNext Ultra II DNA Library Prep Kit for Illumina (New England Biolabs) and sequenced as 40 bp paired-end reads on Illumina NextSeq 500 platform.

### Massively parallel sequencing, data processing and visualisation

For cChIPseq, paired-end reads were aligned to concatenated mouse (mm10) and spike-in human (hg19) genomes using Bowtie 2^63^ with the ‘–no-mixed’ and ‘–no-discordant’ options specified. Reads that were mapped more than once were discarded, followed by removal of PCR duplicates using Sambamba^64^.

For cChIP-seq visualization and annotation of genomic regions, mouse reads were randomly downsampled based on the spike-in ratio in each sample^11^. Individual replicates (n=3) were compared using multiBamSummary and plotCorrelation functions from deepTools (version 3.1.1)^65^, confirming a high degree of correlation (Pearson’s correlation coefficient > 0.9). Normalised replicates were pooled for downstream analysis. Genome-coverage tracks for visualization on the UCSC genome browser^66^ were generated using the pileup function from MACS2^67^ for cChIPseq and genomeCoverageBed from BEDtools (v2.17.0)^68^ for cnRNAseq.

Heatmap and metaplot analysis for cChIPseq was performed using computeMatrix and plotProfile and plotHeatmap functions from deepTools (v.3.1.1)^65^, looking at read density at transcription start sties of all genes. Intervals of interest were annotated with read counts from merged replicates, using a custom-made Perl script utilising SAMtools (v1.7)^69^.

### RNA Fluorescence in situ hybridisation protocol and imaging

smRNA-FISH was carried as described in detail prevously^21^. Briefly, cells were trypsinised and fixed in 3.7% formaldehyde in suspension and then incubated in 70% ethanol at 4C for at least 1h. Cells were then labelled in 2x SSC, 10% formamide and 20% dextran sulfate at 37C overnight with a suspension of 48 20-22nt probes (Stellaris) designed to be evenly distributed across exons or introns of the target transcript. Cells were then spun down and washed multiple times to ensure low nonspecific signal. The cells were then incubated with DAPI to label DNA and Agglutinin-Alexa488 to label cell membranes. The cell suspension was mixed 1:1 with Vectashield H-1000 (Vectorlabs), distributed as a monolayer on glass slides, and covered with microscopy-grade glass coverslips. Images were acquired using the same microscopy setup as described for live-cell transcription imaging except a 2x magnifying lens was used resulting in 91.5nm camera pixel size. In order to estimate mRNA half-life, transcription initiation was blocked with triptolide (500 nM) for 4h and mean numbers of transcripts in cell population were estimated using smRNA-FISH as described above. The experiment was performed in 3 biological replicates. A mono-exponential decay was assumed to represent the mRNA degradation rates upon transcription block and was used to extract mRNA-half-life.

### Live-cell transcription imaging

Transcription was imaged using an Olympus IX83 system fitted with humidified chamber with carbon dioxide-atmosphere at 37C. The microscope was operated through CellSens software and was equipped with a ×63 1.4-NA oil objective lens and a 1,200 × 1,200 px^2^ sCMOS camera (Photometrics). Additional magnifying 1.6x lens was used in front of the camera resulting in final pixel size of 114.4 nm. To image transcription, cells were plated on gelatinised 8-well microscopy µ-slide (IBIDI) 5h in advance of imaging. 1h before imaging the medium was changed to mESC medium with Fluorobrite DMEM instead of Phenol Red DMEM without or with 500 µM auxin in neighbouring wells of the imaging chamber. The imaging conditions were: 20 images at 0.7 µm z-step interval per frame, 8h total duration with 4 min time interval. 20% 490nm exciting light and 70ms camera exposure time were used. A minimum of n=3 biological replicates of untreated and IAA-treated cells were recorded except for *Hspg2* where 2 replicates were acquired.

### Identification of active transcription sites in movies

Individual 3D time-course movies were inspected for cells where there was appearance of transiently accumulating nuclear MCP-GFP signal corresponding to nascent transcription. These cells were cut out and saved as single cell movies. For foci intensity read-out the following protocol was used: first, the custom made ImageJ/FiJi script removed the background with rolling ball algorithm (5 px radius) leaving only punctate MCP-GFP signal. Next, 3D Objects Counter^70^ was applied to individual 3D time-frames to identify active transcription sites in 3D (15 intensity threshold and 10-250 voxel objects). Resulting individual .csv files contained spot volume, intensity, and center of gravity in 3D in individual time-frames. The extracted 3D positions were used to confirm correct spot identification in raw movies.

In order to create time-course fluorescence intensity trajectories for individual active transcription sites (see Figure 1C for examples) a custom made R script was used. Overall, the script uses previously obtained .csv files with MCP-GFP spot detected in individual time-frames to extract the fluorescence intensity of the nascent transcription site and creates a combined fluorescence intensity trajectory. In the case of multiple spots detected in a single time-frame, e.g. when multiple active transcription sites or individual rapidly diffusing pre-mRNAs were identified within the same cell and time-frame *t,* the algorithm follows the spot with the shortest 3D Euclidean distance to the spot it already followed in a preceding time-frame *t-1*. If multiple spots were identified in the first time-frame of the movie (*t*=1), the spot to follow as the transcription site was assigned manually. Every single-cell movie and preliminary trajectory were manually inspected.

These preliminary fluorescence intensity trajectories were then corrected for photobleaching in a following manner. MCP-GFP expressing cells were imaged with identical imaging protocol to the one used for live-cell transcription imaging. The constant background intensity value was measured outside the cells and subtracted from every image. Resulting cell images containing only fluorescence signal were thresholded in 3D using “Huang” settings and total cellular MCP-GFP signal intensity in each time-frame was measured. The resulting normalized GFP photobleaching curve representing 3 biological replicates was approximated with a single exponential fit used next to correct active transcription site fluorescence trajectories through multiplying the extracted transcription site intensity in every time frame *i* by 1/exp(−0.05\**i*), hence accounting for GFP photobleaching during the measurements (Supplementary Figure S3B). Finally, corrected time-course fluorescence trajectories of single active transcription sites were plotted and manually inspected through comparing to raw single-cell movies. A minimum of 250 cells were imaged per biological replicate of which a fraction underwent transcription as judged by MCP-GFP signal accumulation.

### Single pre-mRNA intensity estimation

In order to capture individual pre-mRNAs reliably a slightly altered imaging protocol was used. Briefly live cells were imaged in 3D using 20 images at 0.7µm z-interval with 70ms camera exposure time (same conditions as used for live-cell transcription imaging), however, a 2x magnifying lens was used (image pixel size 91.5nm), and resulted in less light arriving at the camera (0.5723 +/− 0.006 (n=3 measurements)), and this value was taken into account in single pre-mRNA fluorescence intensity calculation (see below). Exciting light was set at 3x the exciting light intensity used for live-cell transcription imaging of active transcription sites. For example, 490nm excitation was set to 83% instead of 20% which corresponded to 3x higher 490nm excitation intensity as evident from calibration curve acquired with varying 490nm excitation intensity and constant camera exposure time (Supplementary Figure 3C). Candidate single pre-mRNA foci were detected using 3D Objects Counter^70^ after subtracting the background with rolling ball algorithm twice (radius = 10px). Foci were identified in i) 2D maximal projections of 3D images for high confidence identification, and ii) in raw 3D images for actual identification. Foci appearing in both approaches were used further. In order to filter out much brighter spots representing active transcription sites a maximal volume threshold of 58.6×10^-3^ µm^3^ was applied, the remaining foci were confirmed to be nuclear and were assumed to represent single pre-mRNAs. Their intensity was measured and was further multiplied by 1/0.5723 = 1.747 (GFP intensity difference originating from using 2x instead of 1.6x magnifying lens, see above) and divided by 3 (to account for 3x the 490nm excitation intensity used in comparison to actual live-cell transcription imaging protocol). Final single pre-mRNA intensity distributions followed normal distribution with mean(standard deviation) of 323(134), 335(115), 330(116) for *Zic2*, *E2f6*, and *Hspg2*, respectively (Supplementary Figure S3D).

### Analysis of transcription parameters from fluorescence tracks

Transcription ON-periods were directly identified in fluorescence trajectories of individual active transcription sites as signal intensity maxima using a custom made algorithm in R. Briefly, the algorithm starts through loading an individual trajectory and uses inflection point identification in order to attribute individual data-points with local maxima or minima with three degrees of strength based on how pronounced they are with respect to surrounding data-points. Time points where no spot was identified (intensity equal to 0) were automatically set as global minima. The algorithm then plots the trajectories with overlaid candidate preliminary maxima and minima for user inspection. Furthermore, every maximum identified in a fluorescence track was inspected. In order to identify an ON-period, a given maximum is assigned a single nearest preceding minimum because every transcription ON-period begins when the fluorescence signal of active transcription site sharply increases and ends when it reaches a maximum. In case no minimum preceding the scrutinized maximum is immediately found while another local maximum is reached, this “intermediate maximum” is discarded from the analysis and the global minimum search continues until one is found. When a minimum-maximum pair is matched, fluorescence signal intensity in time-frames preceding the maximum is investigated in order to identify the true end of the ON-period. This relies on the fact that the ON-period ends when the fluorescence signal ceases to rapidly increase. However, often the global maximum is identified several time-frames away due to fluorescence signal fluctuation and the noisy nature of these data. Therefore, in order to identify the time-frame best representing the end of an ON-period the algorithm studies the local relationship of the identified maximum with five preceding frames and resets its position to the time-frame where the steep signal increase stops. The final minimum-maximum pair represents an individual ON-period. The following parameters are extracted from each ON-period: i) duration time (in minutes), ii) amplitude (in transcripts after converting the arbitrary units of fluorescence into single mRNAs), and iii) RNA Pol II reinitiation rate or time interval between initiating polymerases. In order to approximate the reinitiation rate, fluorescence signal between respective minimum and maximum within ON-period is approximated using a linear fit where its slope represents the speed of transcript production within an ON-period. The rate of polymerase reinitiation can only be estimated for ON-periods greater than 1 transcript. Additionally, due to the 4 min interval used in time-course measurements this analysis could only be reliably carried out for ON-periods with amplitudes exceeding 2.5 transcripts (examples are presented in Supplementary Figure S3E, F).

### Measurements of the fraction of time a promoter spends in the Permissive state

Permissive-periods were identified from live-cell transcription trajectories as consecutive periods in which ON-periods occurred within 60 minutes of each other. Periods outside of permissive-periods were considered OFF-periods. To account for the OFF-periods that occurred in cells lacking detectable ON-periods during the entire 8h-long trajectory, we assumed that each cell contained on average 3 alleles, consistent with ESCs spending a large fraction of their cell cycle in S-phase. Assuming alleles are regulated independently of each other (as shown previously^21^) the number of alleles in a permissive-period per cell should follow a negative binomial distribution of cells with 3, 2, 1, or 0 alleles being transcriptionally permissive during the movie. Therefore, the fraction of the cells where no alleles were transcriptionally active was measured (such cells occurred in 8h-long movies at 36.4(5)%, 40(5)%, and 10(3)% for *Zic2*, *E2f6* and *Hspg2*, respectively) and used to simulate a negative binomial distribution of alleles transcriptionally permissive during the movie recapitulating the abundance of the cells with 0 alleles that are permissive to transcription (or all 3 alleles are in OFF-state). These distributions (obtained at negative binomial probabilities of 0.284, 0.260, and 0.545 for *Zic2*, *E2f6,* and *Hspg2*) were then used to account for all the alleles in cell population that remained in the OFF-state throughout the entire duration of 8h-long movie for UNT cells. For the IAA treated condition the following values were obtained: cells with 0 alleles permissive to transcription comprised 11(2)%, 18(1)%, and 9(9)%, for *Zic2*, *E2f6* and *Hspg2*, and the respective probabilities used to simulate negative binomial distributions were 0.65, 0.4355, and 0.555. Lastly, the total duration of permissive-periods for all the alleles was summed and divided by total measurement time (integrated time spent in OFF- and permissive-periods) to obtain a fraction of time promoter spends in permissive-period.

### RNA-FISH in cell colonies

The cells were plated on 8-well IBIDI µ-well chamber (IBIDI) 12, 24, and 48h prior to fixation with 3% parafolmaldehyde. Then the cells were permeabilized at 37C using 0.5% Triton X-100 for 20 min. RNA-FISH proceeded overnight as described above. Colonies of varying size were manually identified and imaged in 3D using the microscope parameters described above. A custom made Fiji/ImageJ script was used to manually segment the colonies and cut out maximal projections of individual cells that were then subject to transcript counting using ThunderSTORM^71^ as described previously^21^.

### Stochastic simulations of transcript-per-cell distributions

The permissive-period of the promoter was characterized and the number of ON-periods and time between them was measured (Supplementary Figure S4A, B). First, we simulated permissive-periods assuming the number of ON-periods follows a Poisson distribution. We further expected that our 8h-long microscopy measurements may not be able to reliably capture all ON-periods within a permissive-period and instead can be expected to randomly sample it (Supplementary Figure S4C, cartoon). In order to interpret correctly this experimentally assessed number of ON-periods per movie (Supplementary Figure S4B) and account for the fact that our microscopy measurement may capture only a part of permissive-period, we sampled the simulated permissive periods knowing the time interval between ON-periods (Supplementary Figure S4A) using an 8h-long theoretical measurement sliding window recapitulating our microscopy measurements. The number of ON-periods were then counted within that sliding window resulting in the number of ON-periods that would be captured experimentally. We then performed this simulation for a range of hypothetical Poisson-distributed numbers of ON-periods per theoretical permissive-period (Supplementary Figure S4C) and found a value of ON-periods per permissive-period (Supplementary Figure S4D) resulting in a distribution best matching those obtained experimentally (presented in Supplementary Figure S4B). This was done through finding a minimum of 3^rd^-degree polynomial fit (Supplementary Figure S4C). This strategy allowed us interpret the experimentally measured number of ON-periods in 8h-long microscopy experiments and revealed that number of ON-periods per movie measured experimentally for *Zic2* and *E2f6* (Supplementary Figure S4A) corresponded to Poisson-distributed ON-periods per permissive period with mean of 8.95 and 9.33, respectively (Supplementary Figure S4D).

In order to simulate dynamic transcription of *Zic2* and *E2f6* we directly measured ON-period amplitudes (Figure 2B), time intervals between ON-periods (Supplementary Figure S4A), and we inferred number of ON-periods per permissive-period (Supplementary Figure S4D). Hence, the simulation of the Polycomb target gene was assumed to have 3 promoter states, i.e. an allele may either be in i) an OFF-period (no transcription allowed) or ii) in a permissive-period where transcription may take place during iii) ON-periods with known amplitudes (Figure 2B) approximated with a mixed negative binomial and Poisson model which was then used to randomly draw number of transcripts produced per ON-period. Similarly, time intervals between ON-periods, were determined by the number of ON-periods per Permissive-period drawn from Poisson distributions (Supplementary Figure S4D). We simulated individual cells over a period of two 12h-long cell cycles to allow transcript accumulation. For simplicity each cell was assumed to have on average 3 alleles (due to relatively short G1-phase in mESCs). Cell cycles were followed by a cell division resulting in random halving the transcript number with 0.5 probability (Supplementary Figure S4F). Each allele was attributed either OFF-or permissive-period based on a fixed probability p_O>P_ parameter; each allele drew either of the two and was allowed to repeat the draw once at the onset of the 2^nd^ simulated cell cycle. Lastly, a third cell cycle of randomly varying duration (0-12h) was run to desynchronize the cells. At the end the simulation was stopped and simulated cells containing transcripts accumulated over the full course of simulation were subject to transcript degradation with exponentially distributed survival probability dependent on individual transcript age estimated experimentally (Supplementary Figure S4E) such that “old” transcripts were more probable to be degraded. Lastly, a transcript-per-cell distribution was obtained having simulated 500 cells.

Simulations were run for a range of p_O>P_ probabilities and the most similar to the experimental mRNA/cell distribution was identified through minimizing the sum-difference between experimental smRNA-FISH and simulated transcript-per-cell distributions (Supplementary Figure S4G). Using this approach, we identified p_O>P_ values for *Zic2* and *E2f6* in their UNT state. In order to simulate derepression following PRC1 depletion we added an extra step to account for IAA treatment leading to transcript increase: we simulated transcription for an extra 4h (*Zic2*) and 2.5h (*E2f6* as we previously noted it derepresses with a delay^21^) where the p_O>P_ probability value was now increased while all the other transcription parameters were fixed and set to the same values for UNT simulations (ON-period amplitude distribution, duration between ON-periods, and number of ON-periods per Permissive-period). We varied the number of alleles attributed to the cells to account for their different cell cycle stage (cells contained now either 2, 3, or 4 alleles in OFF-or permissive-periods). This strategy allowed us to test if increased p_O>P_ probability can explain the shift in transcript-per-cell distributions following PRC1 depletion (Figure 2D, 3F). Through testing a range of p_O>P_ values we identified those that recapitulated experimental IAA-treated smRNA-FISH distributions best (Supplementary Figure S4G, bottom).

### Single Particle Tracking

Cells were plated a day before on gelatinized microscopy dishes with No. 1.5 (MatTek, #P35G-1.5-14-C). On the day of measurement the cells were labelled using 100nM PA-JF549-Halo (kind gift of L. Lavis and J. Grimm)^72^ for 15 min at 37C, followed by washing 3 times with live cell imaging medium where regular DMEM was replaced with Fluorobrite DMEM (Thermo Fisher Scientific). After 30 min the cells were washed twice before the live cell imaging medium was supplemented with 30mM HEPES.

Single particle tracking was performed using the previously described system^60^ equipped with EMCCD camera (Andor, resulting pixel size 96nm), 100x 1.4NA objective (Olympus) with objective collar and heated stage maintaining it at 37C, laser module (iChrome MLE MultiLaser engine, Toptica Photonics), and translational module (ASI) carrying the fiber optics output used to adjust the beam position between epi- and HiLO-illumination. For imaging at high camera rate 22mW of 561nm laser excitation was used with varied 405nm excitation to maintain fluorescent signals at low density. 4000 15ms frames were acquired per measurement, at least 20 independent measurements containing typically several cells each were acquired per biological replicate. A minimum n=3 biological replicates were acquired for each protein studied.

For stable binding time measurements, after photoactivating sufficient molecules with 405nm laser, long camera exposure time was used (0.5s) and images were acquired with 0.1mW 561nm excitation at different rates for different proteins to adequately address their stable binding: 600 frames at 2Hz for CDK7-HT, HT-CDK9, NELF-B-HT, HT-TFIIB, and HT-NC2β, while for HT-RPB1, HT-TBP and HT-TAF11, HT-dTAG-TAF1, and HT-MED14 200 frames at 0.33Hz was used. Experiments were acquired in a minimum n=3 biological replicates with a minimum of five movies each and an independent H2B-HT control was measured alongside each replicate to correct for photobleaching (see below).

### Single Particle Tracking analysis

Single molecule signals were localized with subpixel resolution using stormtracker software^73^ running in MATLAB (MathWorks) performing elliptical Gaussian point spread function fit to each single molecule signals detected based on fixed intensity threshold (same for all the experiments). Molecule localizations, when appearing in consecutive frames within 8 pixel distance (768 nm) were merged to form tracks (single frame gap was permitted to account for molecule blinking). The resulting track files were converted to an evalSPT format recognized by Spot-ON online analysis tool^42^ used to determine molecule bound fraction through assuming each protein exists in three dynamics states: freely diffusing, slowly diffusing, and bound. The following Spot-ON parameters were applied: 0.01µm length distribution bin width, 10 time points, 10 jumps permitted, maximum jump length of 5.05µm. A localization error of 40nm was assumed, Z correction of 0.7µm, cumulative density function fitting with three iterations. Diffusion coefficient D was estimated as previously described^73^ for tracks that spanned minimum 4 frames. The resulting Log10(D) distributions were fitted with mix of two Gaussians (mixtools R package) and mobility fractions corresponded to their weights.

### Stable molecule binding time estimation

To estimate stable protein molecule binding times bound molecules were localized using stormtracker^73^. Subsequently, tracks representing bound molecules were created after identifying signals appearing in consecutive time frames no further away than 192 nm (2 Hz measurements) or 288 nm (0.33 Hz measurements). The distribution of track lengths of stably bound molecules was fit to estimate apparent dwell times τ:

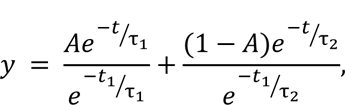

where y denotes fraction of molecules remaining bound at time t, A represents fraction of the first component of molecules with dwell time τ_1_ while τ_2_ is usually longer and represents dwell time of the second component extracted to estimate stable binding time (see below). The first time-point is represented by t_1_. Each biological replicate was accompanied by a separate H2B-HT control measurement representing permanently bound molecules. H2B apparent binding time τ_H2B_ was assumed to be limited solely by dye photobleaching and exceeded that of any measured protein τ_dwell_. Final corrected protein binding time was defined as follows:

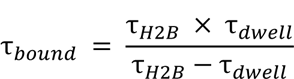

## Acknowledgments

We would like to thank members of the Klose lab for fruitful discussions and for critical reading of the manuscript. We thank Paula Dobrinic, Krzysztof Kus, and Wojciech Siwek for continued input during the project. We thank Johnathan Chubb and Laszlo Tora for insightful scientific discussions. We are grateful to Amanda Williams at the Department of Zoology, Oxford, for sequencing support on the NextSeq 500. We thank Edouard Bertrand for sharing MS2×128 construct and Martin Houlard for providing the T7-SCC1 cells. Work in the Klose lab is supported by the Wellcome Trust (209400/Z/17/Z) and the European Research Council (681440). J.R.K was supported by the Oxford-Wolfson Marriott Graduate Scholarship.

## Contributions

A.T.S. and R.J.K. conceived the project and wrote the paper with contributions from all co-authors. A.T.S. performed most of the experiments, data analysis, and visualisation. E.D performed genomics experiments with analyses. J.R.K. carried out biochemical experiments and contributed to refining the course of the project.

---

**Supplementary Figure S1.**
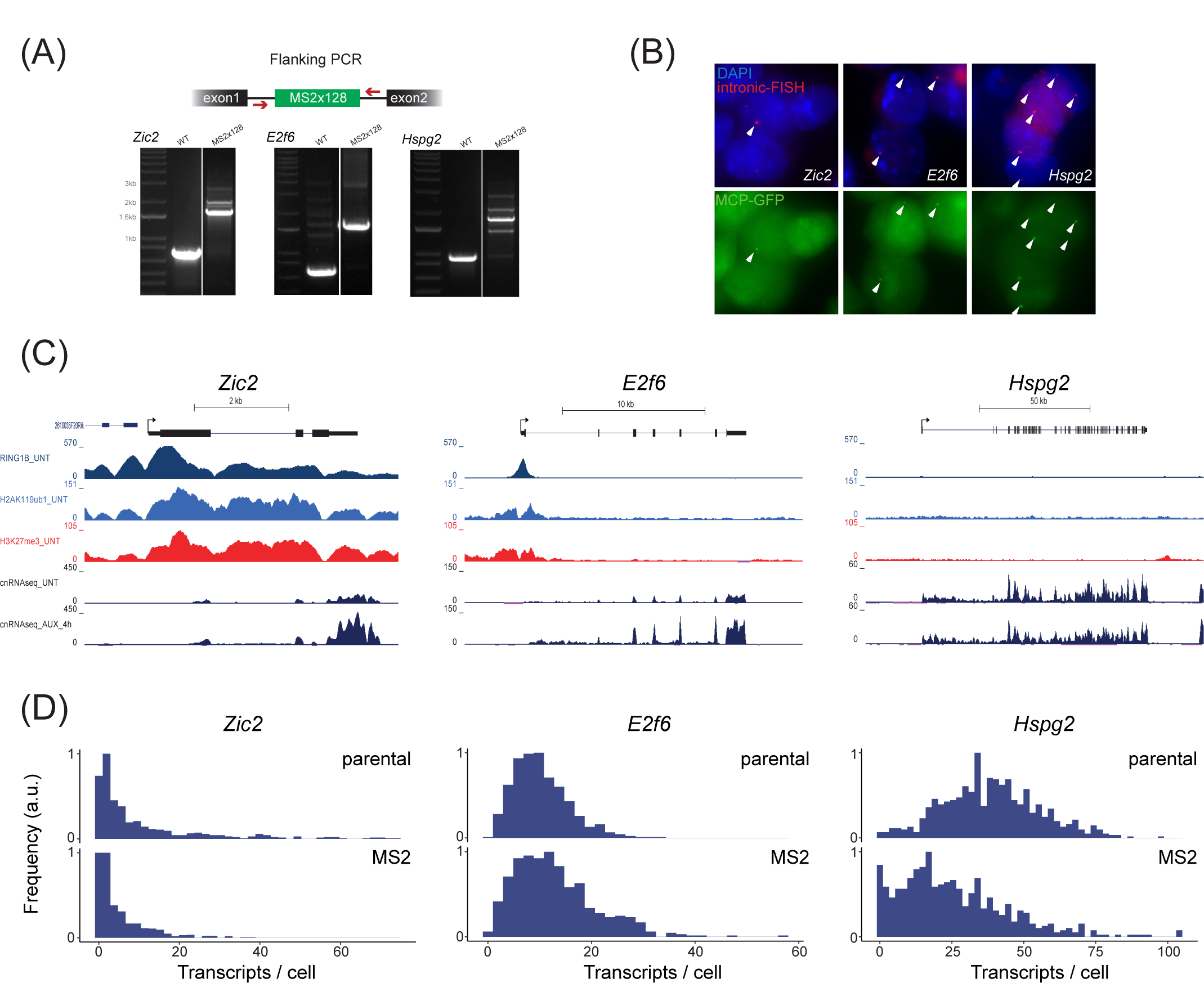
Characterisation of live-cell transcription imaging in ESCs. A) Validation that the MS2×128 array is appropriately inserted into the first intron of the corresponding gene. Top: a schematic illustrating the PRC screening strategy. Bottom: PCR results for *Zic2*, *E2f6*, and *Hspg2*. B) Images of intronic RNA-FISH (red) and focalized MCP-GFP signal (green) indicating that MCP-GFP accumulates at sites where intronic RNA sequences for *Zic2*, *E2f6*, and *Hspg2* are identified. Nuclei are labelled with 4′,6-diamidino-2-phenylindole (DAPI, blue). C) Genomic ChIP-seq snapshots for *Zic2*, *E2f6*, and *Hspg2* illustrating signal for Ring1B-AID, H2AK119ub1, and H3K27me3. Nuclear RNA-seq signal before and after 4h Ring1B-AID depletion is also shown^21^. D) smRNA-FISH analysis of transcript-per-cell distributions for parental (MCP-GFP expressing) and MS2×128 array-containing cell lines.

**Supplementary Figure S2.**
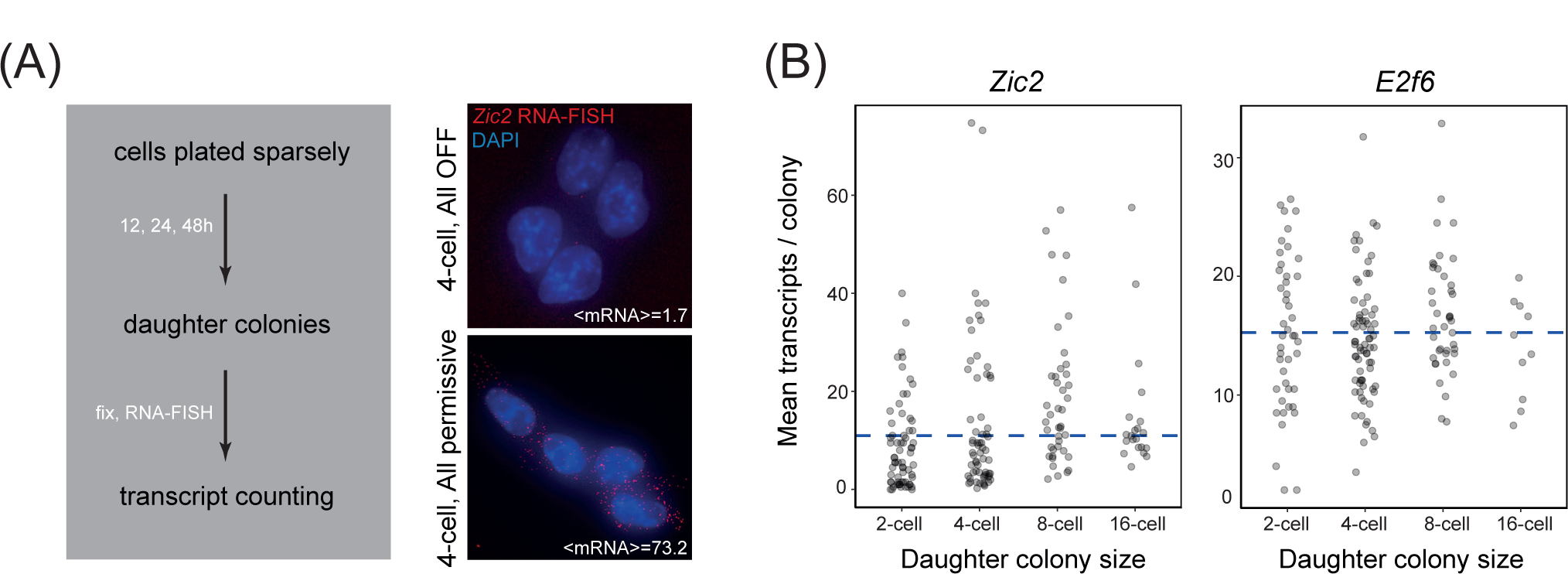
Testing heritability of transcription activity of Polycomb-targets across cell divisions. A) A strategy to assess the number of transcripts-per-cell for Polycomb target genes between monoclonal daughter cells (grey box, left). Right: examples of smRNA-FISH images of 4-cell colonies with all cells having or all cells lacking *Zic2* transcripts. This shows that the expression state of Polycomb target genes can be heritably retained across cell divisions. B) Mean number of Polycomb target gene transcripts per colony vs. colony size. Individual dots represent measurements for single monoclonal colonies. The blue dashed line represents the mean number of transcripts-per-cell in all colonies measured. Note, highly- or non-expressing colonies are still found in 4-cell colonies (2 cell divisions) indicating the respective state has been maintained across cell divisions.

**Supplementary Figure S3.**
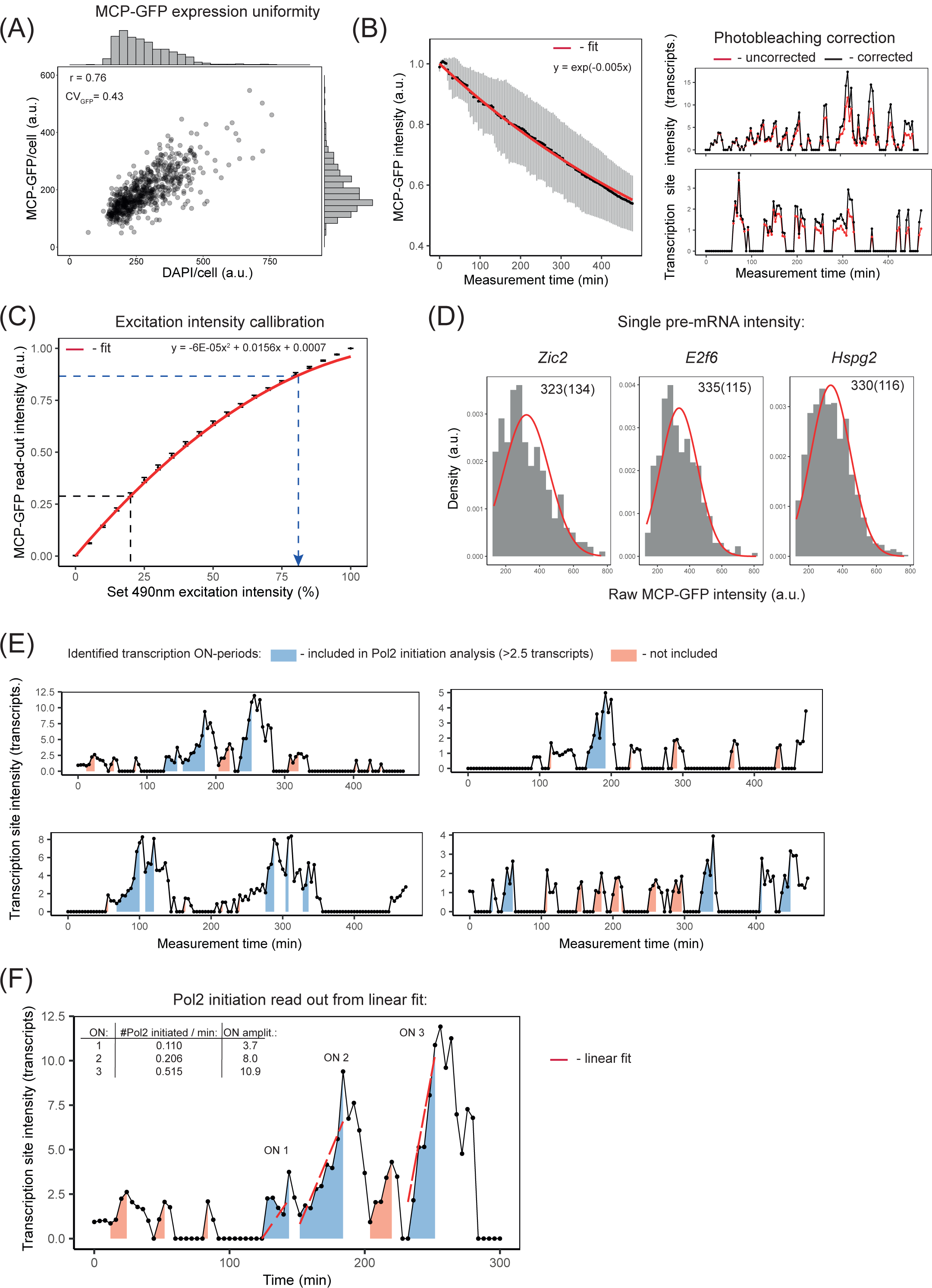
Characterisation of live-cell transcription imaging with single-transcript sensitivity and ON-period analysis. A) MCP-GFP expression is uniform across the cell population correlated with DNA content (DAPI signal). B) (Left panel) Measurements of GFP photobleaching (grey datapoints) over a full time-course of live-cell-imaging approximated with an exponential decay (red line) that was used to correct fluorescence intensity in time-course transcription trajectories. (Right panel) Examples of the effect of this correction are presented on the right. C) To measure the intensity of single pre-mRNAs containing 128 MS2 aptamers, imaging was performed using a higher 490nm excitation intensity. The curve quantifies MCP-GFP intensity (y-axis) in response to varying 490nm excitation levels. The blue dashed lines represent values used for live-cell transcription imaging and for single pre-mRNA intensity quantification (dashed line with arrow-head). This curve informed us of the 490nm intensity that excites GFP at 3x the value used in our live-cell transcription measurements. D) Histograms of single pre-mRNA intensities recalculated in values corresponding to live-cell transcription measurements for *Zic2*, *E2f6*, and *Hspg2*. The red line represents a Gaussian fit with mean and standard deviation values indicated above. These values allowed us to recalculate fluorescence intensity units in order to attribute transcript numbers based on fluorescence intensity at the transcription site. E) Examples of live-cell transcription trajectories with identified ON-periods indicated in blue or orange depending on whether they were taken into account during RNA Pol II reinitiation rate estimations or not. All ON-periods were taken into account in amplitude and duration analysis. F) An example of a live-cell transcription trajectory with three ON-periods (in blue) with their amplitudes and RNA PolII reinitiation rates (from linear fits, red dashed lines) indicated.

**Supplementary Figure S4.**
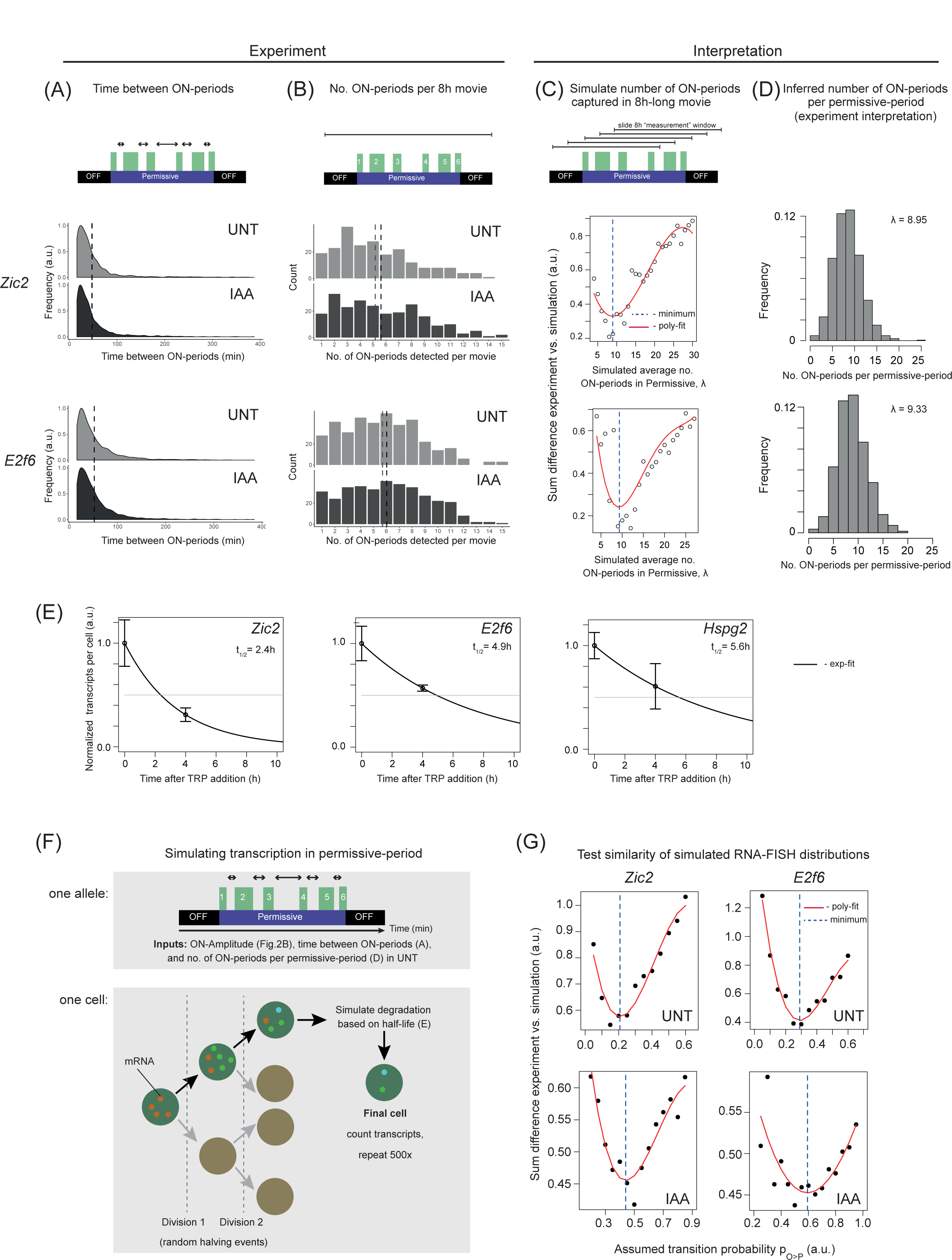
Stochastic simulations of transcription to obtain transcript-per-cell distributions and estimate transition probability from OFF-to Permissive-states for Polycomb targets. A) Density plots of time intervals between ON-periods (indicated as arrows in the cartoon) directly measured from live-cell transcription imaging trajectories for both Polycomb targets *Zic2* (top) and *E2f6* (bottom) for untreated (UNT) and PRC1-depleted (IAA) conditions. Dashed vertical lines represent mean values. ON-, permissive-, and OFF-periods are indicated in the cartoon in green, purple, and black, respectively. B) Histograms of number of ON-periods detected per 8h live-cell transcription movie (indicated in the cartoon as blunt-end horizontal line). Dashed vertical lines represent mean values. C) In order to interpret the detected number of ON-periods per 8h movie and infer the number of ON-periods in a permissive period, the permissive periods were simulated with varying mean Poisson-distributed number of ON-periods (λ, x-axis) and “sampled” using a “sliding” 8h window to represent the experimental measurement (blunt-end horizontal line in the cartoon). The sum difference between the resulting distribution and experimental distribution (presented in B) was calculated (y-axis). The red line represents 3^rd^-degree polynomial fit and its minimum (vertical dashed line) represented the mean number of ON-periods expected to produce most similar distribution of captured ON-periods per 8h measurement window. Plots for *Zic2* (top) and *E2f6* (bottom) are shown. D) Histograms of inferred mean number of ON-periods per permissive period for *Zic2* (top) and *E2f6* (bottom). E) Estimates of transcript half-lives for *Zic2*, *E2f6*, and *Hspg2*. Data-points represent normalized mean number of transcripts in untreated (t=0) and after 4h of triptolide (TRP) treatment obtained by smRNA-FISH. Solid black lines represent exponential fits. Horizontal grey lines represent half of the mean transcript number detected in untreated sample. The intersection between black and grey lines indicates transcript half-life. F) A cartoon illustrating the strategy to simulate transcription of Polycomb-target genes. (top) At an individual allele level every parameter of transcription necessary to simulate the permissive-state is quantified or inferred: ON-period amplitude (in transcripts), time between ON-periods, and number of ON-periods in a permissive state. (bottom) Cells were assumed to have on average 3 alleles, and were allowed two full cell cycles followed by cell divisions leading to random halving of the transcript numbers. Single cells were simulated leading to transcript accumulation. Once produced, transcripts were attributed a date-of-birth which was used at the end of the simulation to degrade transcripts based on mRNA half-life. This procedure was repeated 500 times to produce simulated single-cell distribution of transcripts-per-cell. G) The procedure described in (F) was repeated using a range of probabilities of transitioning between OFF- and permissive-states (p_O>P_) to produce simulated transcript-per-cell distributions that were then compared to smRNA-FISH experimental data and the most similar were identified by the minimum in 3^rd^ degree polynomial fit (red line) indicated as vertical blue line for *Zic2* (left) and *E2f6* (right) in untreated (UNT) or PRC1-depleted (IAA) conditions.

**Supplementary Figure S5.**
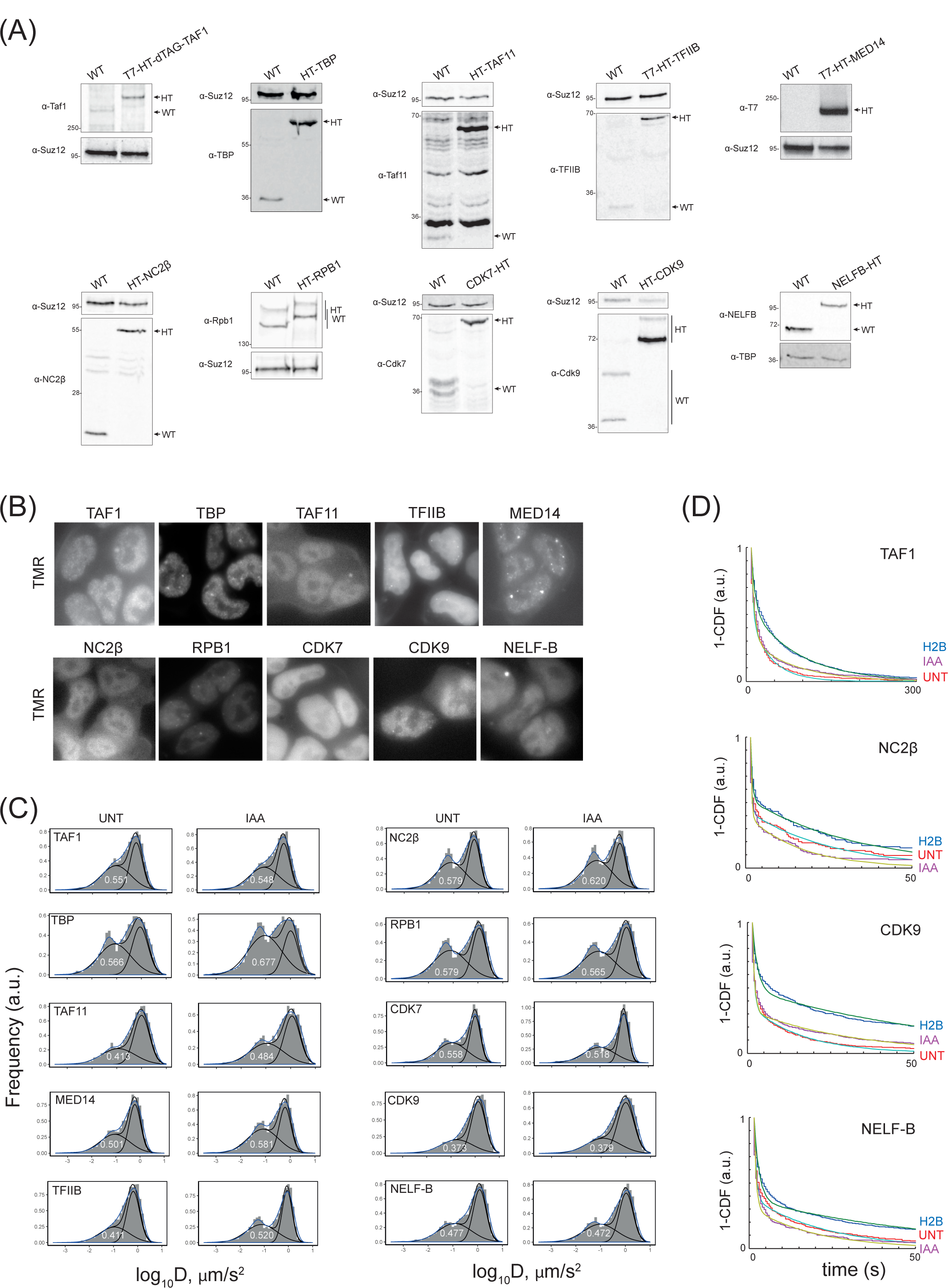
Extended data to single-particle tracking of transcription regulators. A) Western blot analysis of endogenously HALO-tagged factors comparing the signals in wild type and tagged lines. Antibodies and molecular weight markers (in kilodaltons (kDa)) are indicated on the left, wild type (WT) and HALO-Tag (HT) protein bands are indicated on the right with arrows. B) Microscopy validation of the HALO-Tag expression in lines with endogenously tagged proteins. HALO-Tag-proteins were visualised using TMR-HALO ligand. All proteins localized to the nucleus. C) Examples of representative biological replicates of histograms of log10(D) calculated from single-particle tracking data acquired at high camera frame rate, obtained for the panel of transcription regulators with (UNT) and without PRC1 (IAA). Black solid lines represent a mixed two-Gaussian fit (to account for immobile and mobile fractions) with indicated value representing immobile portion of molecules. Blue solid line represents histogram density. D) Examples of 1-CDF plots representing single molecule binding times acquired at low camera frame rate. Average stable binding time is extracted from bi-exponential fits indicated in the plots. Examples of data acquired with (UNT, red line) and without PRC1 (IAA, purple line) together with respective H2B-HT (blue). The latter represents a stable binding control used to correct photobleaching.

**Supplementary Figure S6.**
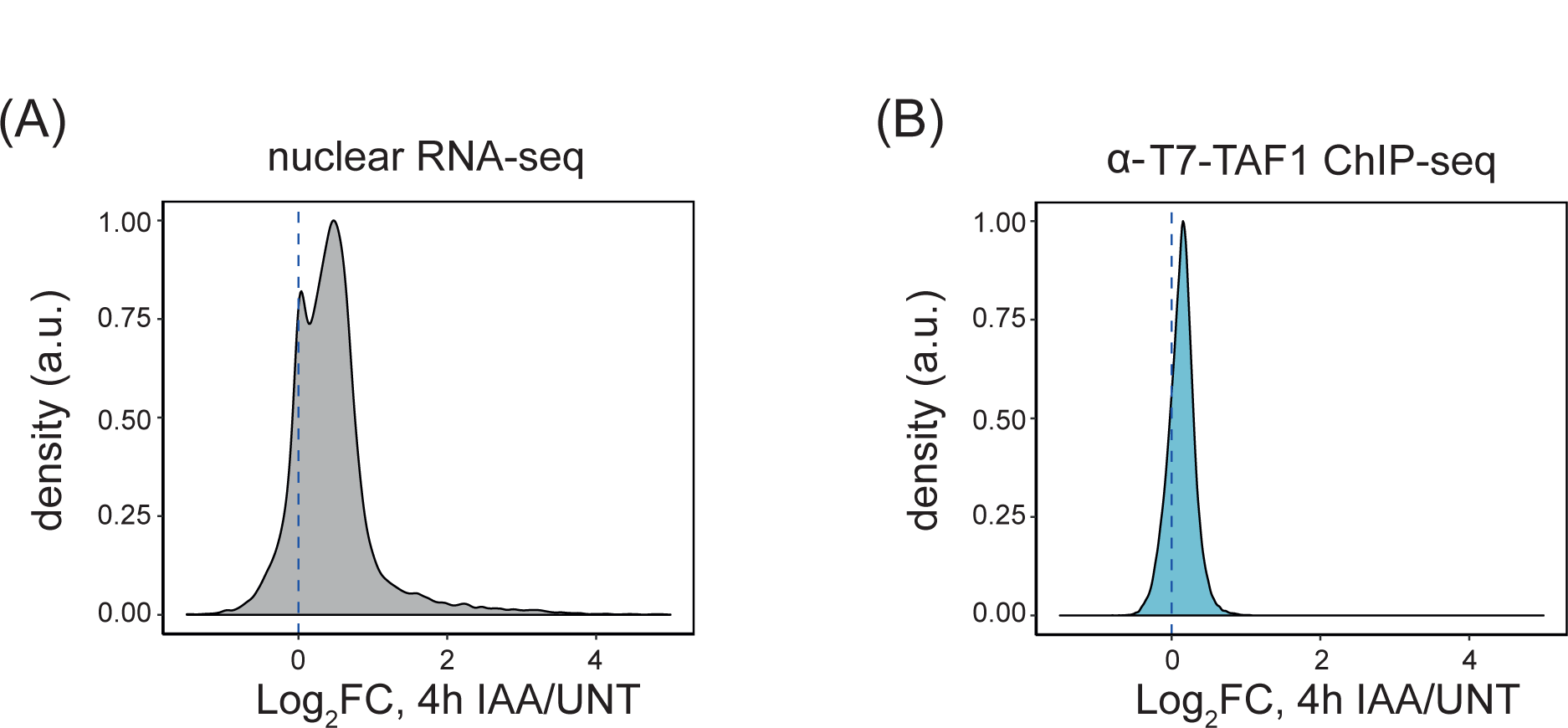
Genome-wide effects on gene expression exerted by Polycomb-depletion. A) RNA-seq log2 fold change (LFC) density plot for all genes comparing untreated to 4h PRC1 depletion gene expression levels^21^. Vertical dashed line represents no change. B) Log2FC density plot comparing integrated TAF1 ChIP-seq signal within a 1kb window centered at TSS. All genes are included. Vertical dashed line represents no change.

